# Elucidating the Role of Primary and Secondary Sphere Zn^2+^ Ligands in the Cyanobacterial CO_2_ Uptake Complex NDH-1_4_: The Essentiality of Arginine in Zinc Coordination and Catalysis

**DOI:** 10.1101/2024.01.20.576359

**Authors:** Ross M. Walker, Minquan Zhang, Robert L. Burnap

## Abstract

The uptake of inorganic carbon in cyanobacteria is facilitated by an energetically intensive CO_2_-concentrating mechanism (CCM). Specialized Type-1 NDH complexes function as a part of this mechanism to couple photosynthetic energy generated by redox reactions of the electron transport chain (ETC) to CO_2_ hydration. This active site of CO_2_ hydration incorporates an arginine side chain as a Zn ligand, diverging from the typical histidine and/or cysteine residues found in standard CAs. In this study, we focused on mutating three amino acids in the active site of the constitutively expressed NDH-1_4_ CO_2_ hydration complex in *Synechococcus* sp. PCC7942: CupB-R91, which acts as a zinc ligand, and CupB-E95 and CupB-H89, both of which are in close interaction with the arginine ligand. These mutations aimed to explore how they affect the unusual metal ligation by CupB-R91 and potentially influence the unusual catalytic process. The most severe defects in activity among the targeted residues are due to a substitution of CupB-R91 and the ionically interacting E95 since both proved essential for the structural stability of the CupB protein. On the other hand, CupB-H89 mutations show a range of catalytic phenotypes indicating a role of this residue in the catalytic mechanism of CO_2_-hydration, but no evidence was obtained for aberrant carbonic anhydrase activity that would have indicated uncoupling of the CO_2_-hydration activity from proton pumping. The results are discussed in terms of possible alternative CO_2_ hydration mechanisms.

## 1. Introduction

Cyanobacteria and other photosynthetic organisms have evolved mechanisms to actively uptake, store, and use inorganic carbon to overcome limitations in the main CO_2_-fixing enzyme, Rubisco. These mechanisms are collectively known as the CO_2_ concentrating mechanism (CCM), which functions by increasing CO_2_ concentration around Rubisco [1-6]. The cyanobacterial CCM can be conceptually divided into two concurrently operating processes: 1.) uptake processes leading to the accumulation of high levels of bicarbonate (HCO_3_^-^) in the cytoplasm and 2.) utilization processes taking the accumulated bicarbonate into the carboxysome microcompartment, which encloses the cellular complement of Rubisco and delivering saturating concentrations of CO_2_ to the Rubisco active sites due to the action of a co-localized carbonic anhydrase. Uptake of inorganic carbon in cyanobacteria occurs via five distinct uptake protein complexes, each contributing to the accumulation of HCO_3_^-^ in the cytoplasm. Bicarbonate is directly acquired from the environment through the three identified types of cytoplasmic membrane transport proteins: SbtA, BicA, and BCT1 [1-6].. The SbtA and BicA act as Na^+^-HCO_3_^-^ symports and thus actively transport across the cytoplasmic membrane HCO_3_^-^ using the inwardly directed Na^+^ electrochemical gradient. The BCT1 complex is an ABC-type transporter and uses the energy of ATP hydrolysis to drive HCO_3_^-^ into the cytoplasm. Meanwhile, CO_2_ can be actively converted to HCO_3_^-^ within the cytoplasm by specialized forms of the NAD(P)H dehydrogenase type-1 (NDH-1) complex. NDH-1 are members of the large Complex I family of quinone-reducing, proton-pumping bioenergetic complexes that, in cyanobacteria, exist in multiple forms, including those involved in CO_2_ uptake. There are two paralogous NDH-1 CO_2_ uptake systems, NDH-1_3_ and NDH-1_4_, which catalyze high affinity, low flux and low affinity, high flux CO_2_ uptake, respectively. Each uses the redox energy-transducing machinery to actively convert CO_2_ to HCO_3_^-^ (CO_2_ hydration), although the nature of this coupling remains obscure. The CO_2_ hydration complexes allow both the direct uptake of CO_2_ diffusing into the cell and the re-capture of CO_2_ lost from the carboxysome. Collectively, the HCO_3_^-^ transport and CO_2_ uptake activities in cyanobacteria produce a high internal HCO_3_^-^ pool in the range of 10-40 mM [7-9]. Bicarbonate, being ionic, is a form of C_i_ that is about 1000-fold less permeable to lipid membranes than the uncharged CO_2_ molecule and, therefore, is more readily accumulated within the cytosol. The far-from-equilibrium accumulation of HCO_3_^-^ has been graphically demonstrated by the heterologous expression of human carbonic anhydrase within the cytosol of *Synechococcus elgonatus* sp. PCC 7942 (hereafter, *Syn*7942) [10]. This led to the continuous dissipation of the HCO_3_^-^ pool due to the rapid equilibration between bicarbonate and CO_2_ and the consequent high-flux leakage of CO_2_ from the cell. High cytoplasmic HCO_3_^-^ is used by the other major part of the cyanobacterial CCM, namely the carboxysome [11-15]. The carboxysome contains the entire cellular complement of Rubisco, along with a CA. High cytoplasmic HCO_3_^-^ levels drive a massive diffusive flow into the carboxysome, where the CA efficiently converts the HCO_3_^-^ into CO_2,_ thereby effectively saturating the Rubisco active site with CO_2_ and minimizing the competing and wasteful photorespiratory reaction with O_2_ [16].

The mechanisms of energy conversion and CO_2_ uptake in NDH-1_3/4_ complexes remain poorly understood. Pioneering genetic studies identified high C_i_-requiring mutants with lesions in genes encoding NDH-1 proteins [17-21]. These studies also revealed the presence of a truly novel class of proteins that help catalyze CO_2_ hydration (CupA and CupB that are also essential for CO_2_ uptake. The CupA/B proteins form their own class of proteins in the Pfam database with no sequence or protein-fold similarity to any known carbonic hydrases [22]. The CupA and CupB proteins are subunits of the NDH-1_3_ and NDH-1_4_ complexes, respectively. The complexes can be differentiated via regulation and their affinities for CO_2_ during the uptake process [20, 23-26]. The low-affinity system, NDH-1_4_, is constitutively expressed, whereas the high-affinity system, NDH-1_3,_ is induced by low CO_2_ [20, 21, 24, 26]. The multiplicity of NDH-1 complexes includes the form, designated NDH-1_1/2_ of the mediating respiratory electron flow and cyclic electron flow (CEF), which appears to be the dominant form of the complex present in the cyanobacterial cell [27], although the high affinity NDH-1_3_ complex becomes comparatively abundant under high light conditions [28]. This functional diversification with the cyanobacteria is manifested by the existence of multiple isoforms of the NdhD and NdhF proteins. There are typically five paralogs of the NdhD protein, NdhD1-5, and three paralogous forms of the NdhF protein, NdhF1,3, and 4, which are also present in cyanobacterial genomes. All isoforms of the NDH-1 complex share a common ‘core’ module that consists of the hydrophilic ‘peripheral arm,’ which coordinates iron-sulfur centers that mediate electron transfer from ferredoxin to plastoquinone. This common core shared amongst the multiple NDH-1 isoforms also contains approximately half of the portion of the transmembrane subunits involved in proton pumping. The different forms of complex expressed in the cells can distinguished by their complement of NdhD and NdhF proteins, as well as the presence or absence of the CupA/B proteins and various smaller subunits. The NDH-1_1/2_ complexes contain the common core plus the NdhD_1/2_ and NdhF_1_. The CO_2_ uptake NDH-1 complexes contain the common core plus subcomplexes consisting of [NdhD_3_-NdhF_3_-CupA/S] and the [NdhD_4_-NdhF_4_-CupB].

Structural analysis using cryo-EM has revealed intact structures of cyanobacterial NDH-1_1/2_ respiratory complexes [30-33], as well as, most recently, the structure of the high-affinity CO_2_ hydration complex NDH-1_3_ [34]. The structures share conserved features found across the large family of Complex I bioenergetic complexes and also, finally, resolve the donor of high energy electrons as ferredoxin (Fd) rather than NAD(P)H, as found in the respiratory complexes of many heterotrophs [35]. The putative catalytic site CO_2_ hydration in the NDH-1_3_ complex is situated at the interface of the CupA protein and the transmembrane antiporter domain of the NdhF and was shown to have a Zn ion (**Fig. 1**). The Zn-containing active site of the CupA/B proteins of the CO_2_ uptake NDH-1_3/4_ complexes is inferred to have CA activity somehow coupled to the redox and/or the proton-pumping activity of the complex [1, 36]. This activity is unique among known CAs because it is coupled to energy transduction in NDH-1_3/4_ complexes in a manner that drives the CO_2_ hydration reaction far from equilibrium and in the direction of bicarbonate formation. All other CAs are energetically uncoupled and simply mediate the formation of an equilibrium state between the reactants and products. It is, therefore, not surprising that the active site structure in the NDH-1_3/4_ complex does not closely resemble known CAs. Most notably, the presence of an arginine side chain as a Zn ligand (**Fig. 1**) is unlike anything previously described for metal ligands of the various classes of CAs^1^. Canonical CAs have three amino acid ligands to the metal, usually a combination of histidine and/or cysteine residues. A fourth water ligand acts as a substrate and undergoes deprotonation during the catalysis, with the resultant Zn-hydroxide anion engaging in nucleophilic attack upon incoming CO_2_ molecule, yielding the bicarbonate product of the anhydrase reaction [38, 39]. The CupA/CupB proteins have only two amino acid ligands: an arginine side chain (CupA-R135/CupB-R91) ligating via its guanidinium moiety and a histidine side (CupA-H130/CupB-H86) chain providing ligation by its imidazole nitrogen. The remaining Zn coordination positions may be occupied by water molecules, although the resolution of the *Thermosynechococcus* NDH-1_3_ cryo-EM structure does not allow a definitive assignment. Besides the novel metal coordination, other structural features of the CupA/B active sites are likewise intriguing. These include the highly conserved presence of a second arginine sidechain adjacent to the Zn and a tyrosine proposed by an acceptor of protons released from the substrate water early in the catalytic cycle [34]. We have recently produced a homology model for the NDH-1_4_ complex of the genetically amenable *Syn*7942 [29]. The CupB-R91 residue is modeled to coordinate active site Zn corresponding to CupA-R135 in the paralogous CupA protein of the high-affinity Cup system in the *Thermosynechococcus* NDH-1_3_ structure. The second amino acid ligand to the Zn ion is modeled to be CupB-His86, which has been previously shown to be essential for CupB protein accumulation [29]. The arginine and histidine Zn ligands are conserved across the CupA/B protein family, consistent with their crucial roles in coordinating the active site metal center. Here, we describe the effects of mutations involving the replacement of amino acids carried out at the unique zinc-binding residue Arg91 and the two adjacent second-sphere residues, His89 and Glu95, which interact with the primary ligand Arg91 through their side chains potentially modulating the unusual metal ligation by CupB-R91 and potentially influencing the proton transfer reactions integral to the catalytic process.

**Figure 1.**
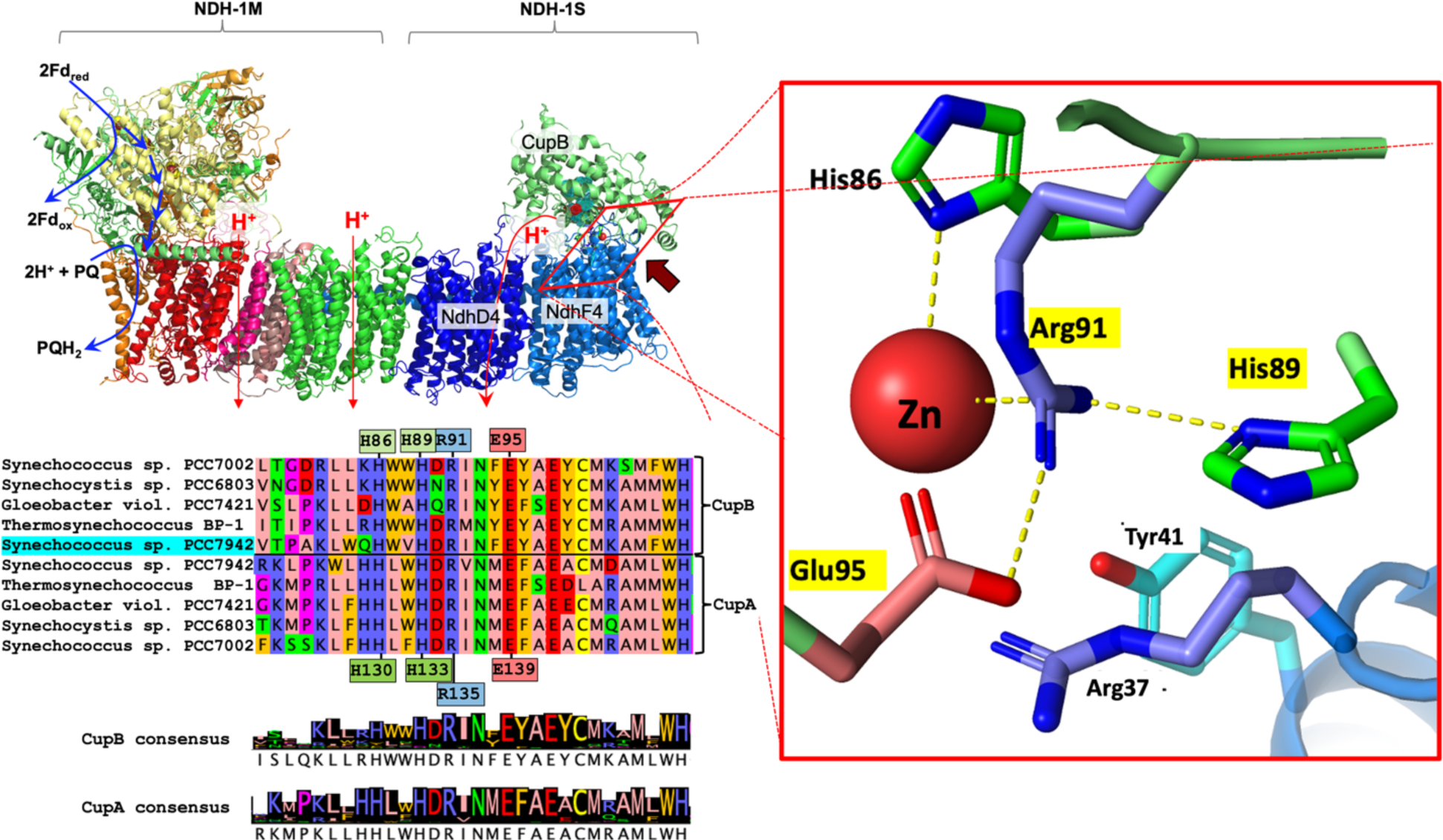
The active site of CO_2_ hydration. Multiple sequence alignment (MSA) of Cup proteins A and B from a relevant selection of cyanobacterial genomes (inset, lower left). Correspondence between rows indicates homology according to MSA (ClustalW) analysis described previously [29]. Six species are shown as an excerpt from a larger alignment of CupA and CupB protein sequences identified from 157 cyanobacteria strains. The excerpted sequences correspond to *Synechococcus* sp. PCC7002 (Syn7002), *Synechocystis* sp. PCC6803 (*Syn6*803), *Gloeobacter violaceus* sp. PCC7421, *Thermosynechococcus elongatus* BP-1, *Synechococcus* sp. PCC7942 (*Syn*7942). The active site (the featured region of the sequences) shows high conservation as indicated by the consensus logos of the CupA and CupB protein subfamilies. Homology modeling of the putative CupB active site based on the CupA subunit of *Thermosynechococcus* from the NDH-1_3_ published coordinates (PDB 6TJV) developed previously [29] as shown (top left and right side, respectively). The mutationally targeted residues H89, R91, and E95 of CupB are indicated with yellow highlighted labels in the close-up of the Zn-containing active site (right side). The highly conserved residues around the putative active site of CupB and NdhF4 are shared by their respective homologs, CupA and NdhF3. The model shows two residues, CupB-H86 and CupB-R91, acting as direct ligands to the Zn ion, consistent with the homologous side chains of the CupA protein of the *Thermosynechococcus* NDH-1_3_ structure.

## 2. Materials and methods

### 2.1 Cyanobacterial strains and growth conditions

The wild-type strain of *Synechococcus* sp. PCC 7942 (hereafter *Syn*7942) and its genetically modified derivatives were maintained in liquid and solid agar BG-11 medium [40] supplemented with 10mM HEPES-NaOH pH 8.0 amended with appropriate antibiotics. Experimental cultures were grown in the absence of antibiotics in 1 L Roux bottles, continuously bubbled (approximately 150 mL min^-1^) with air enriched with 3% or 5% CO_2_ at 30°C with a light intensity of ∼200 μmol (photons) m^-2^ s^-1^ using fluorescent lights.

### 2.2 Construction and expression of CupB mutants

The generation of experimental strains began with the construction of a strain that had both *cupA* and *cupB* deleted from the genome [9]. This recipient strain was maintained in BG-11 media containing 25 μg mL^-1^ chloramphenicol (Cm) and 10 μg mL^-1^ kanamycin (Km) and is designated *ΔcupA/ΔcupB* (originally designated *ΔXΔY* [9]). The reintroduction of wild-type and mutant variants of the *cupB* gene was accomplished as described previously [29] by gene insertion into the polylinker region of the pSE4 shuttle vector, which is capable of replication in both *Escherichia coli* and *Syn*7942 [41]. Due to the presence of abundant nitrate in the growth medium, the *nir* promoter driving clone gene expression is active under all growth conditions and is, therefore, an effective promoter for the constitutive expression of *cupB* [42, 43]. Site-directed mutant gene fragments were commercially synthesized (GenScript Inc).

### 2.3 Isolation and immunoblot analysis of thylakoid membrane and soluble fractions

Cells were harvested from cultures grown under 3% CO_2_ to an OD_750_ ∼0.8 via centrifugation at 4000 x *g* for 6 minutes at room temperature and resuspended to 1 mg Chl mL^-1^ in thylakoid breakage buffer (TBB: 20mM HEPES, 100mM NaCl, 10% glycerol (w/v)). The concentrated cells in TBB were broken by agitation with 0.1 mm zirconium beads (1:1). Cell homogenate was separated into thylakoid membrane (TM) and soluble components by centrifugation at 40,000 x*g* for 40 minutes. The blue cytoplasmic fraction was removed and stored in TBB, and the TM fraction was resuspended in TM-Suspension Buffer (TSB: 50 mM MES-NaOH, 10% (v/v) glycerol, 1.2 M Betaine (monohydrous, Sigma), 5 mM MgCl_2_, 20 mM CaCl_2_) and adjusted to ∼ 0.5 – 1 mg Chl mL^-1^. Isolated TM fractions from CupB mutants were mixed 1:1 with sample buffer (125 mM Tris-HCl, 20% glycerol, 2% sodium lauryl sulfate (SDS), 0.05% bromophenol blue, 0.1 M dithiothreitol for SDS-PAGE separation and electrotransferred to a polyvinylidene difluoride membrane (Immobilon-FL; Merck Millipore) and detected by an antibody raised against synthetic peptides of *Synechococcus elongatus* PCC sp. 7942 CupB (PhytoAB-Antibody).

### 2.5 CO_2_ uptake assays for measuring C_i_ affinity of whole cells

Cells grown under ∼200 μmol (photons) m^-2^ s^-1^ of light and bubbled with CO_2_-enriched air (5%). were harvested via centrifugation at 4000 x *g* for 6 minutes at room temperature. Cells are then washed and resuspended in pH 7.0, Ci-free, Na^+^-free 50mM BTP buffer as described previously [29]. Samples were exposed to ∼400 μmol (photons) m^-2^ s^-1^ illumination in the sample chamber of a Clark-type electrode until samples approached the O_2_ exchange compensation point. Following this, the uptake of C_i_ was estimated based on the rates of O_2_ evolution as C_i_ was titrated into the chamber in a stepwise fashion. Potassium bicarbonate (KHCO_3_^-^) was added in the presence of 25 μg mL^-1^ carbonic anhydrase (CAII from bovine, Sigma, USA) to the pH 7.0 BTP buffered cells in the electrode chamber. Under these conditions, most C_i_ will take the form of CO_2_.

### 2.6 Measurement of carbonic anhydrase (CA) activity

Carbonic anhydrase activity was measured as the rate of change in pH following the addition of ice-cold CO_2_ saturated water (prepared by bubbling ice-cold water with 100% CO_2_ for at least 1 hour prior to measurements) to a 20mM Tris-HSO_4_^-^ assay buffer (pH 8.3) in the presence and absence of freshly isolated thylakoid membranes (TMs) as described above. Assays were performed on ice using a pH electrode inserted into 6 mL of assay buffer stirred. A TM sample corresponding to 20 µg Chl was added, and after equilibration of temperature and pH, 4 mL of ice-cold CO_2_-saturated water was pipetted into the assay buffer. Ice cold water was added in place of a TM sample as a negative control to measure the uncatalyzed rate of CO_2_ hydration. Carbonic anhydrase II (CAII: Sigma-Aldrich, C2624) was added in place of a TM sample as a positive control to measure the catalyzed rate of CO_2_ hydration. One Wilbur-Anderson unit (WAU) of activity is defined as (*T*_0_ − *T*)/*T*, where *T*_0_ (uncatalyzed reaction) and *T* (catalyzed reaction) are recorded as the time (s) required for the pH to drop from 8.3 to 6.3 after the addition of ice-cold CO_2_ saturated water [44].

## 3. Results

### 3.1. CupB active site mutagenesis

Directed mutagenesis targeting the amino acid residues of the putative catalytic active site of the CupB protein in the *Synechococcus* 7942 NDH-1_4_ complex was conducted based on previous homology modeling [29] based upon the *Thermosynechococcus elongatus* NDH-1_3_ cryo-EM structure [34]. Substitution mutations were introduced at the unusual Zn-ligand, Arg91 and the two second sphere ligands, His89 and Glu95, whose side chains interact with the primary ligand Arg91. All three residues are strictly conserved in both the CupA protein and CupB protein subfamilies, as shown by multiple sequence alignment (MSA) **(Fig. 2)**. The mutants were introduced into the Δ*cupB*/Δ*cupA* strain [9], which entirely lacks CO_2_-hydration capabilities associated with the NDH-1_3/4_ complexes due to the absence of both CupB and CupA proteins. Reintroduction of the wild-type and modified *cupB* genes was achieved using a replicative plasmid [29]. Transformation the Δ*cupB*/Δ*cupA* strain using the plasmid containing the wild-type version of *cupB* is referred to as the WT control (WT-C), although it is not the true wild-type in the sense that it lacks the ability to express the CupA protein. As noted previously, the CO_2_-hydration activity of the NDH-1_4_ complex functions normally with the plasmid-expressed CupB protein enabling the study of CO_2_ uptake activity due exclusively to the WT control CupB protein and its mutant derivatives [29].

**Figure 2.**
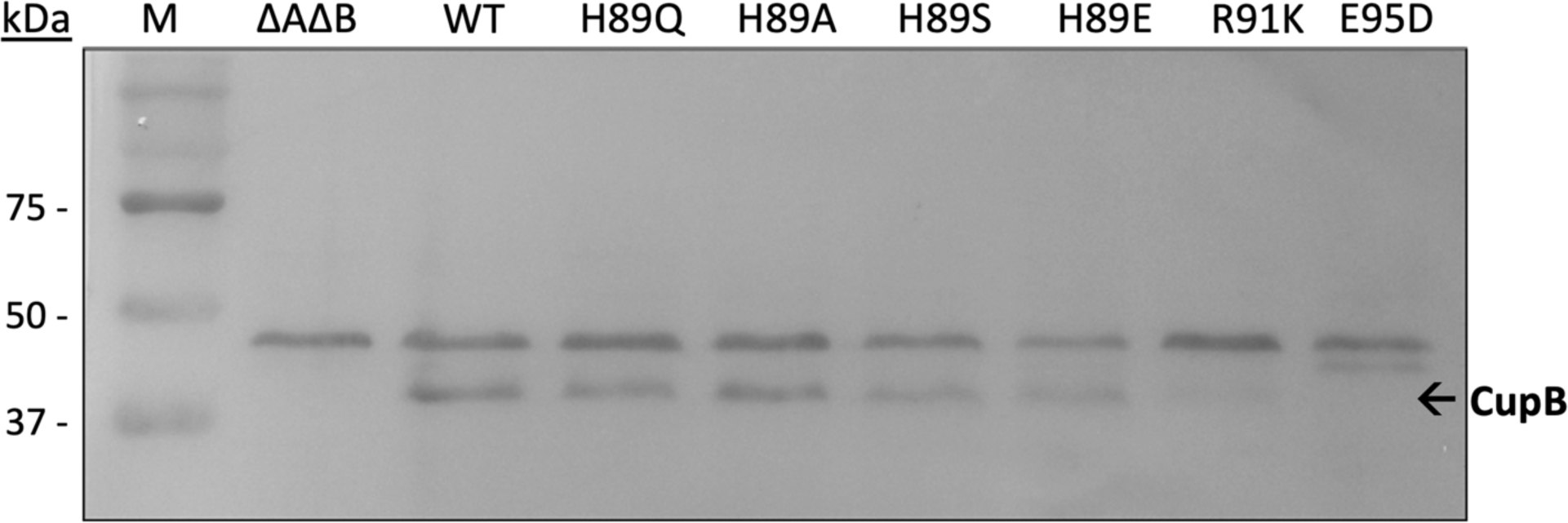
Immunodetection of CupB accumulation in mutants *Syn*7942 strains. Membrane protein isolates of *Syn*7942 strains lacking both CupA and CupB (ΔAΔB), expressing the wild-type CupB protein in the ΔAΔB genetic (WT-C) background and various CupB amino acid substitution mutants are probed with CupB antibody purified from rabbit immune serum. Cells were grown at 3% CO_2_ in BG-11 medium at pH 8.0 and crude thylakoid membranes were isolated as described in Methods. Cytoplasmic fractions were also probed for the presence of CupB but none was detected (Fig S1). After solubilization, polypeptides in the membrane samples were analyzed by SDS-PAGE, electroblotted to a polyvinylidene difluoride (PVDF) membrane, and probed with anti-CupB antibody. CupB is expected to have a mass of 42 kDa.

### 3.2. Contrasting Effects of His89, Arg91, and Glu95 Substitutions on CupB Accumulation

CupB accumulation was analyzed by immunoblot analysis using polyclonal antibodies directed against the CupB protein. The CupB protein is an extrinsic membrane protein bound primarily to NdhF4, but with extensive contacts with NdhD4 transmembrane subunits of the NDH-1_4_ complex (Fig. 1). To evaluate the possibility that mutant CupB variants may not assemble properly with the remainder of membrane embedded NDH-1_4_, cell homogenates were separated into membrane and cytoplasmic fractions for immunoblot analysis. All CupB-His89 mutants accumulate CupB to levels similar to that observed in the wild-type control in the thylakoid membrane fraction, presumably attached to NDH-1_4_ complex (**Fig. 1**). In contrast, all amino acid substitutions at CupB-Arg91 and CupB-Glu95 positions eliminate CupB protein accumulation (**Fig.S1)**. Substituting Arg91 with lysine (CupB-R91K), glutamine (CupB-R91Q), and histidine (CupB-R91H) and the substitutions of Glu95 with glutamine (CupB-E95Q), alanine (CupB-E95A), serine (CupB-E95S), and aspartate (CupB-E95D), led to a stark reduction in the CupB accumulation, to undetectable levels in most instances, except for a marginal presence observed in the CupB-E95Q mutant (**Fig S1A, C**). Since no accumulation of the CupB mutant proteins was observed in the cytoplasm (**Fig. S1**), it appears that the inability to fold properly and/or bind to the remainder of the NDH-1_4_ complex renders the mutant CupB protein susceptible to proteolytic degradation, although more direct evidence for the putative degradation would require further study. Nevertheless, both CupB-Arg91 and CupB-Glu95 are clearly crucial for the stability of the CupB protein. In the context of the structural model (**Fig. 1**), it is postulated that the Arg91 residue, besides serving as a ligand to the catalytic Zn, forms a pivotal ionic or H-bond with Glu95, that underpins the structural stability of CupB folding and ensures a secure attachment to the membrane portion of the complex. At the same time, CupB-His89 is also closely positioned to Arg91, yet mutations at this position do not as strongly affect the accumulation of the protein (**Fig. 2**). Thus, we can conclude that CupB-His89 is not crucial for the structural integrity of CupB, in contrast to the structurally critical CupB-Arg91 and CupB-Glu95 residues. However, as shown in the next section, CupB-His89 is important for the catalytic efficiency of the enzyme, which is consistent with its strict evolutionary conservation.

### 3.3. CupB-H89 plays a catalytic role in CO_2_-uptake

The strains containing CupB-H89 mutations accumulated the CupB protein and were subjected to further analysis. To assess the impacts of point mutations on the CO_2_-dependent O_2_ evolution [29, 45, 46], experiments with whole cells were conducted in a sodium-free buffer, which inhibits the activity of sodium bicarbonate symporters, at pH 7.0 and in the presence of an externally added carbonic anhydrase, thereby focusing the experiment on CO_2_ as the primary source of C_i_ as detailed previously. Figure 3 illustrates the rates of oxygen evolution as a function of C_i_ concentration (primarily CO_2_). The data were analyzed using Michaelis-Menten kinetics to determine the whole cell affinities (K_0.5_) and maximal rates (V_max_) of CO_2_ uptake.

**Figure 3.**
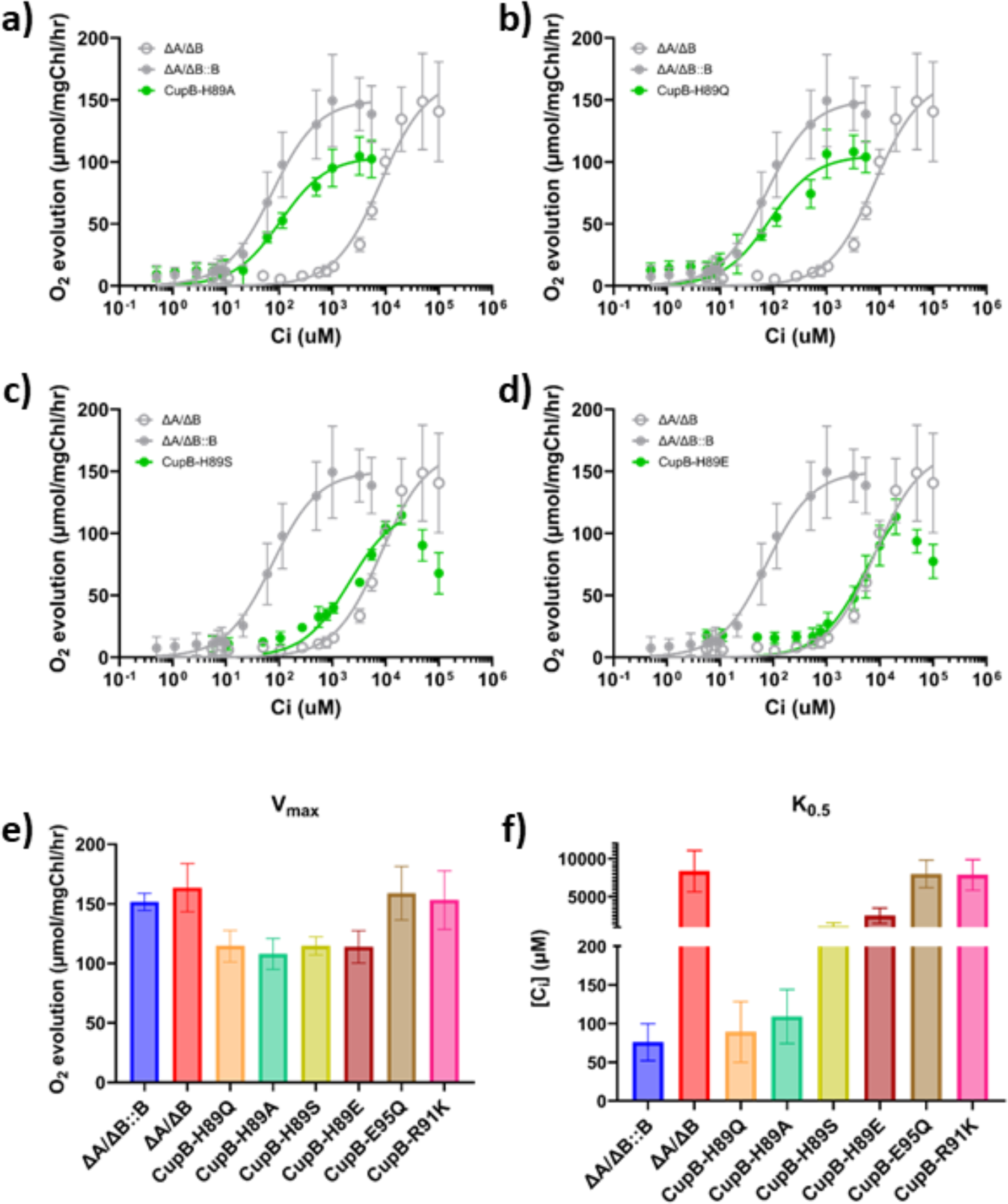
Clark-type electrode measurements of O_2_ evolution as a proxy for CO_2_ uptake. Whole cell response to titration of C_i_ reveals the affinity for CO_2_ in His89 mutants as a function of [C_i_] (a – d). Cells were grown under 5% supplemental CO_2_ to prevent accumulation of low-C_i_ inducible CCM components such as SbtA or NDH-1_3_. Affinity is measured as the [C_i_] at which the cells evolve O_2_ at half their maximum velocity. In this way, the affinity of whole cell mutants can be represented by K_0.5_, similar to the Michaelis-Menten K_m_ unit. n = >3. V_max_ BTP (e) represents the maximum rate of O_2_ evolution measured during the 15-minute affinity assay. K_0.5_ (f) represents the value of [C_i_] which corresponds to half the V_max_. Error bars represent standard deviation from the mean (biological n ≥ 3; technical n ≥ 8).

The CupB-H89 mutations display lower affinities for CO_2_ and lower maximal rates of CO_2_-dependent oxygen evolution, suggesting impacts on the enzyme’s catalytic performance (**Fig. 3a-d**). Whereas cells depending on the wild-type version of the CupB exhibited a K_0.5_ ∼60 μM, consistent with previous results, the CupB-H89E and CupB-H89S mutations severely decreased the concentration of C_i_ required to reach a half-maximum velocity of O_2_ evolution (K_0.5_ ∼ 2000 μM, **Fig 3f**), requiring much higher CO_2_ concentrations to saturate C_i_-dependent photosynthetic oxygen evolution activity. While part of the reduced affinity can be attributed to the reduced abundance of the protein in CupB-H89E and CupB-H89S, significant accumulation occurs, yet its affinity approaches that of *ΔcupB/ΔcupA*, completely lacking CupA and CupB. In contrast, the H89A substitution greatly restored the affinity for CO_2_ (K_0.5_ ∼175 μM). Similarly, the Gln substitution only modestly affected CO_2_ uptake affinity in agreement with previous results [29]. Each His89 mutant also has a lowered V_max_ in comparison to WT-CupB to a surprisingly similar degree (∼25% decrease), considering their varying degrees of impact regarding the affinity. The parent *ΔcupB/ΔcupA* strain lacking CupB exhibits a V_max_ similar to the CupB-WT and higher than any of the H89 mutants, suggesting that the presence of the CupB-H89 mutation has an inhibitory effect on net photosynthesis rates. This cannot be attributed to a decreasing level of PSII activity *per se* since the rates uncoupled of PSII activity using the artificial electron acceptor DCBQ show no significant differences were observed compared to the wild-type control (**Fig. S3 right panel**). In contrast, the CupB-R91K and CupB-E95Q mutants, which fail to accumulate CupB, behave like the *ΔcupB/ΔcupA* double-deletion strain regarding their CO_2_ uptake affinity and V_max_, as expected considering they abolish the accumulation of the protein. It also indicates that the genetic expression system is not responsible for this phenotype. Thus, the mere presence of the mutant CupB-H89 proteins has an inhibitory effect, albeit modest, on maximal rates of CO_2_-coupled oxygen evolution.

### 3.4. No evidence for a mutationally-induced reversible carbonic anhydrase

We had previously hypothesized that CupB-H89Q mutation uncoupled the proton pumping activity, rendering the Zn-centered carbonic anhydrase activity reversible and thus dissipating rather than promoting the formation of high bicarbonate concentrations in the cytoplasm [29]. To investigate this, we conducted carbonic anhydrase activity assays, measuring pH changes in diluted buffers with initially containing saturating concentrations of dissolved CO_2_. Compared to the control using bovine carbonic anhydrase, anhydrase activity in membranes of the WT-control and the CupB-H89Q mutant was very low and, importantly, were equivalent. Thus, this test showed no significant differences in activity between the WT-control and the CupB-H89Q mutant (**Table 1**).

**Table 1.**
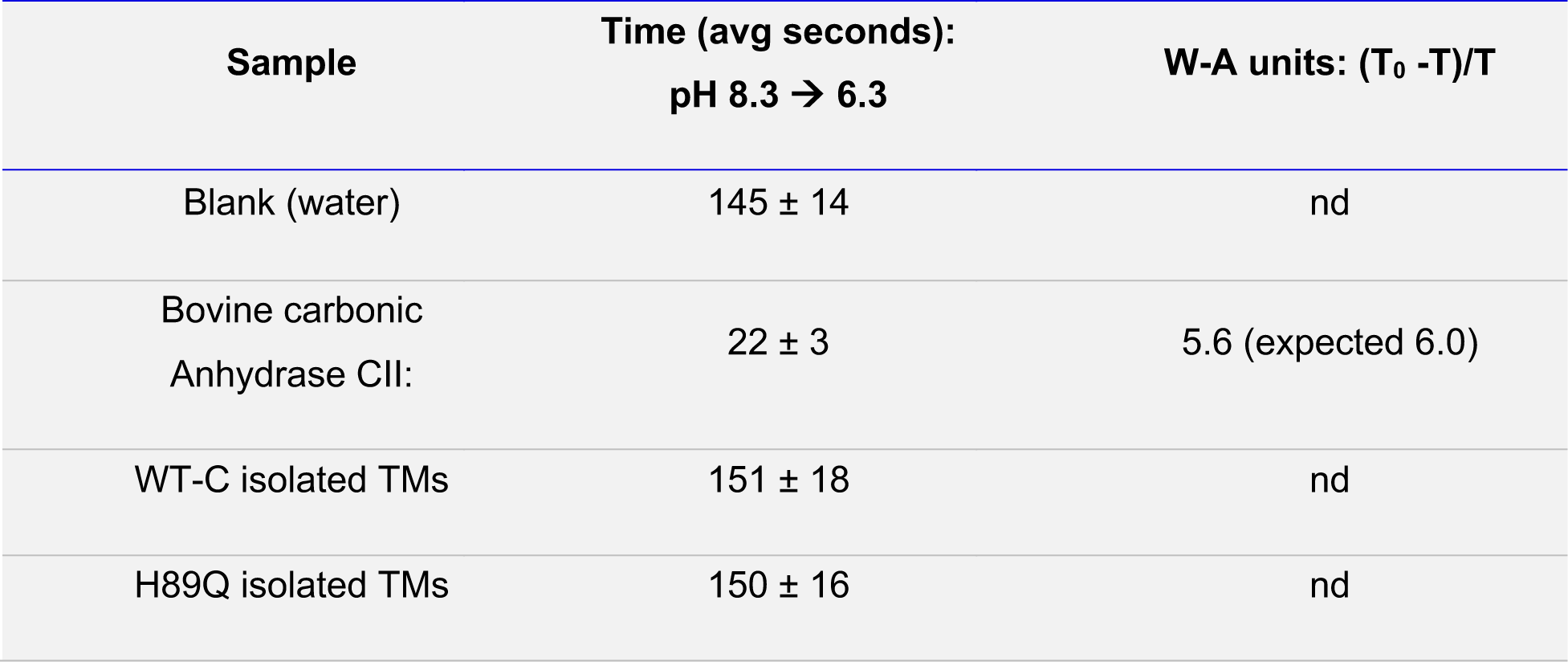
Carbonic anhydrase (CA) activity measured as the rate of change in pH of a weakly buffered solution from 8.3 to 6.3. One WAU is defined as (*T*_0_ − *T*)/*T*, where *T*_0_ (uncatalyzed reaction) and *T* (catalyzed reaction) are recorded as the time (s). Protocol adapted from Millipore Sigma (EC 4.2.1.1).

| Sample | Time (avg seconds):<br>pH 8.3 → 6.3 | W-A units: $(T_0 - T)/T$ |
| --- | --- | --- |
| Blank (water) | 145 ± 14 | nd |
| Bovine carbonic<br>Anhydrase CII: | 22 ± 3 | 5.6 (expected 6.0) |
| WT-C isolated TMs | 151 ± 18 | nd |
| H89Q isolated TMs | 150 ± 16 | nd |

## 4. Discussion

The CO_2_ uptake NDH-1_3/4_ complexes, unique for their powered carbonic anhydrase activity, exhibit an active site with a zinc (Zn) ligand configuration distinct from canonical CAs. Unlike typical CAs that facilitate equilibrium between reactants and products, these complexes drive the CO_2_ hydration reaction towards bicarbonate formation. The atypical active site includes an arginine side chain as a Zn ligand, which is a departure from the usual combination of histidine and/or cysteine residues found in standard CAs (**Fig. 1**). Here we mutationally targeted three amino acid residues in the active site region of NDH-1_4_: the CupB-R91 Zn ligand, and two residues, CupB-E95 and CupB-H89, that interact with the arginine ligand, potentially tuning its ability to engage its unusually metal ligation and possibly mediating the proton transfer reactions that undoubtedly occur during catalysis.

### 4.1. The Arg-Glu dyad critical for CupB stability

The loss of cellular CupB accumulation observed for mutations at CupB-E95 and CupB-Arg91 underscores their pivotal contributions to the enzyme’s structural integrity. Even the relatively conservative mutations of CupB-E95 and CupB-R91 residues proved non-functional (**Figs. 2 & 3**). For example, the conservative CupB-E95D mutation also prevents accumulation of CupB (**Fig. S1**). This indicates that the aspartate side chain, shorter by one methylene group, is unable to engage in a critical interaction with guanidinium of CupB-R91. In the wild-type, there appears to be a close interaction of the wild-type CupB-E95 and CupB-R91, where the distance between the guanidinium and carboxylate is modeled to be less than 3 Å thereby providing a strong stabilizing ionic linkage at this position that appears crucial for structure stability. Also, the CupB-R91K mutation is non-functional. The lysine side chain has essentially the same length as arginine and, therefore, could, in principle, form the critical ionic interaction with CupB-E95. However, CupB, with this conservative substitution also fails to accumulate in cells. The essentiality of the CupB-R91 residue is likely to reflect a complex role since its multifunctional guanidinium moiety enables it to simultaneously interact with the CupB-E95 carboxylate and to offer an lone electron pair as the ligand to the Zn^2+^. In addition, the chemical versatility of the guanidinium moiety may also allow proton transfer functionality. Thus, it seems likely that some or all of these facets of chemical versatility cannot be fulfilled by the Lys side chain. In this regard, it is important to note that mutation of the second amino acid Zn^2+^ ligand, CupB-H86, also prevents accumulation of the CupB protein, pointing to the criticality of Zn^2+^ ion binding for the stabilization of the CupB structure and its interaction with the membrane portion of the complex [29]. Thus, it can be concluded that both the disruption of the Zn^2+^ ligation and the interaction of CupB-E95 and R91 contribute to the accumulation of the CupB protein. In this regard, it is noteworthy that the entire Zn^2+^-binding region of CupB is situated at the interface between the CupB protein and membrane-spanning the NdhF4 subunit. The disruption of Zn ligation and/or the disruption of the intramolecular ionic linkage may perturb the binding interface of the CupB to the membrane complex with the concomitant release from the membrane, rendering it prone to proteolytic degradation as an unfolded protein in the cytoplasm. Alternatively, these interactions may be important early in the biogenesis process as the nascent CupB protein is initially expressed, for example, perturbing proper protein folding and/or Zn-binding, thus rendering the mutant protein susceptible to proteolysis. Interestingly, the computational analysis of the original *Thermosynechococcus* NDH-1_3_ structure suggested that the CupA-R135 Zn ligand, which corresponds to CupB-R91 in our S7942 model, is uncharged due to its interaction with the active site Zn^2+^ ion (*vide infra*) although this remains to be determined experimentally. Even if this were the case, it does not preclude the possibility that a stabilizing ionic interaction is critical during assembly, if not once the fully assembled metalloprotein is formed. Finally, an important significance of the finding that the Cup-H86 and R91 Zn ligand mutants fail to accumulate CupB is that it lends further support to the unprecedented ligation assignments in the cryo-EM structure [34].

### 4.2. CupB-His89 is important for catalytic function

In contrast to mutations of the Arg-Glu dyad, structural integrity is largely maintained in the CupB-H89 mutants, yet the catalytic activity of the enzyme is affected to differing degrees depending on the substitution. The CupB-H89 and the homologous CupA-H133 residue of NDH-1_3_, are highly conserved, suggesting an important functional role for the imidazole side chain at this position. Despite accumulating substantial amounts of the CupB protein, the different CupB-H89 substitutions possess different CO_2_ uptake affinity characteristics, yet all depress steady-state photosynthesis. The glutamine (CupB-H89Q) and alanine (CupB-H89A) substitutions at His89 have modest impacts on CO_2_ uptake affinity. While differing from histidine in terms of hydrogen bonding capacity and titratability, these substitutions do not drastically alter the overall functional environment necessary for catalytic function. However, the glutamate (CupB-H89E) and serine (CupB-H89S) substitutions show a more drastic reduction in CO_2_ affinity. These effects may reflect their inability to support proton transfer or failure to provide an important H-bond. The contrasting behavior of the CupB-H89S and CupB-H89A mutants is particularly interesting. They accumulate the CupB protein, yet the two substitutions have dramatically different impacts on the CO_2_ affinity characteristics of the enzyme, with the serine substitution being more deleterious. These mutants have smaller side chains that potentially allow more water molecules to occupy the space normally taken by the wild-type histidine, and such waters can potentially mediate proton transfer or H-bonding. The intruding waters may provide the necessary H-bonds or provide proton transfer functionality, for example, by a Grotthuss mechanism. A role in proton transfer seems less likely than an H-bonding interaction between CupB-H89 and the Zn^2+^-ligating CupB-R91 residue, as discussed below. Despite their differing impact upon CO_2_ uptake affinity, each of the mutants CupB-H89 depress the maximal rate of CO_2_-dependent photosynthesis and results in PSI acceptor side limitation, suggesting that all the mutations affect steady-state CO_2_ hydration and the provision of bicarbonate to the carboxysome.

### 4.3. No evidence for a reversible carbonic anhydrase reaction

Previous work had shown that several CupB mutants, including the H89Q mutant, exhibited characteristics suggesting that mutant forms of the CupB protein can be more detrimental to cellular [C_i_] status than having no CupB protein whatsoever. The current findings confirm these results, with all the CupB-H89 mutants showing depressed maximal C_i_-dependent oxygen evolution rates and evidence for acceptor side limitation of PS1 (**Fig. S4**). Based on this surprising result, it was proposed that the mutations enabled a latent carbonic anhydrase activity that had become uncoupled from CO_2_ hydration from vectorial proton pumping that normally drives the reaction away from equilibrium towards CO_2_ hydration. The putative carbonic anhydrase activity in these mutants was hypothesized to exert its negative effects on inorganic carbon status by aberrantly dissipating bicarbonate that had been accumulated within the cytoplasm through the action of the other bicarbonate transporters (e.g. SbtA, BicA) of the cell. Here, we tested this hypothesis using isolated membranes but found no evidence for the putative reversible carbonic anhydrase reaction (**Table 1**). Although we cannot exclude the possibility that activity was lost during membrane isolation, we tentatively conclude that either the activity does not exist or that the carbonic anhydrase activity is strictly dependent upon the normal membrane energization processes found *in vivo*.

### 4.4. Models for the modulation of CO_2_ affinity by CupB-H89

The negative result regarding the absence of carbonic anhydrase activity prompted us to consider other models for the directionality of the CO_2_ hydration activity, including the originally proposed mechanism involving the observed CO_2_ channel in the structure [34]. We also considered how the CupB-89 mutant potentially affects catalytic activity in a possible mechanism. Changes in the binding affinity of CO_2_ in the active site in the traditional sense of energetics of the interaction of CO_2_ with the residues of the active site. Instead, changes in affinity are more likely related to the proton-handling and ionization characteristics of the enzyme. If the reaction is indeed chemically analogous to canonical carbonic anhydrases, then the effects on the apparent K_m_ (K_0.5_) in a substrate versus rate experiment are primarily due to impacts the mutations have upon lifetime of the anionic deprotonated state rather than relating to a substrate binding constant for CO_2_. The K_m_ in carbonic anhydrases is remarkably high, in the range of 1-10 mM and higher [47, 48], considering the catalytic efficiency typically displayed by these enzymes. Rather, the very high enzyme efficiency is due to an extraordinary turnover rate (k_cat_ ∼10^6^ s^-1^) that allows it to operate near the diffusion limit with the rates approaching the collision frequency of substrate CO_2_ reaching the active site. Such collisions are productive if they encounter the de-protonated state of the substrate, the Zn-hydroxide species. Accordingly, the deprotonation of substrate H_2_O coordinated to the Zn and subsequent proton transfer away from the active site generally limits the rate of CO_2_ hydration [48, 49]. Thus, the rate of deprotonation and the subsequent lifetime of the resultant Zn-hydroxide species govern the intrinsic catalytic turnover (k_cat_). The longer lifetime of the anionic reactive species also presents as a higher ‘affinity’ for CO_2_: even when CO_2_ concentrations are low, which are conditions where diffusive visits of CO_2_ into the active site are less frequent, productive encounters will still occur between the incoming CO_2_ and deprotonated water. Thus, productive encounters will occur in proportion to the length of the deprotonated state lifetime. Overall, stabilization of the deprotonated Zn-hydroxide enhances the ability of the enzyme to operate under such low [CO_2_] regimes and this can be accomplished by efficient proton shuttling and stabilization away from the Zn-hydroxide intermediate. In human carbonic anhydrase II, histidine 64 (CAII-H64) fulfills the proton acceptor function that stabilizes this hydrolysis intermediate. In this regard, the computational QM/MM MD modeling [34] of the homologous NDH-1_3_ CO_2_ uptake complex indicates that the tyrosine, NdhF3-Y41 (corresponding to NdhF4-Y41, **Figs. 1 and Fig. S5**), fulfills the role of primary proton acceptor of the predicted water deprotonation reaction. Like CAII-H64, the putative Tyr proton acceptor in the NDH-1_3/4_ complex faces the Zn-coordinated substrate water, and intervening water molecules are proposed to mediate proton transfer. The computational model of the NDH-1_3_ active site predicts pK_a_ 6-7 for NdhF3-Y41 and facile Grotthuss transfer via two intervening water molecules stabilizing the predicted reactive Zn-hydroxide [34]. This initial Tyr proton acceptor is in an exposed position situated at the mouth of the active site cavity leading to the aqueous soluble phase of the cytoplasm (NdhF4-Y41 and NdhF3-Y41). In contrast to this, CupB-H89 in NDH-1_4_ and the corresponding CupA-H133 in NDH-1_3_ are buried in the interior of the protein adjacent to the primary Arg Zn ligand. Therefore, if CupB-H89 is stabilizing the Zn-hydroxide to promote high catalytic efficiency, it seems unlikely to be doing so by acting as a secondary proton acceptor in a proton relay starting with NdhF4-Y41 given the interceding NdhF4-R37. However, the imidazole N of H89 appears to act as an H-bond acceptor from the NH2 group of CupB-R91. Since CupB-R91 also coordinates the Zn^2+^ ion, a more likely role for CupB-H89 is to position and potentially tune the Lewis acidity catalytic metal, hence the hydroxide intermediate’s lifetime.

Considering that NDH-1_3/4_ complexes power hydration CO_2_ using their redox and proton-pumping activity, it is likely that key reaction steps are very different compared to typical carbonic anhydrases. This is consistent with the very different Zn-ligation and different constellations of side chains in the active site of the NDH-1_3/4_ complex compared to carbonic anhydrases. Thus, while the NDH-1_3/4_ complex may share the basic feature of employing a Zn ion to activate the bound substrate water, other steps after the nucleophilic attack on incoming CO_2_ may be fundamentally different. Two mechanisms for CO_2_ hydration have recently been proposed. The first involves gated CO_2_ transport based on the observation in the structure of a CO_2_-conducting channel surrounded by hydrophobic and bulky residues that lead from the luminal side of the membrane to the Zn-binding active site [34]. It is proposed that the channel opens and closes depending on the ion-pair conformations in NdhF3/NdhF4 subunit, and this gating provides directionality to the CO_2_ hydration reaction presumably by sterically preventing the reverse reaction [34]. Ion-pair interactions in the NdhF3/NdhF4 subunit are part of a putative energy-transducing charge-relay process proposed to be transmitted through antiporter-like subunits and are responsible for proton pumping across the membrane [50]. The charge-relay process coupled to proton pumping has combined support from computational simulation and experimental studies [51], although its coupling to gas channel gating remains to be tested. An alternative proposed mechanism is essentially a product removal hypothesis where the proton released during CO_2_ hydration is fed into the proton pumping activity of the antiporter-like subunits, and thereby, the hydration reaction is driven by product (H^+^) as discussed [29] (**Fig. S4**) previously. Again, the putative energy-transducing charge-relay process is proposed to be transmitted through antiporter-like subunits, but the proton transport activity itself is hypothetically responsible for coupling the CO_2_ hydration and the bioenergetics of the NDH-1 complex. This model was based upon the phenotype of the CupB-H89Q mutation that suggested a defect in coupling the CO_2_ hydration reaction to proton pumping, rendering the reaction reversible (see Supplemental Figure S. However, we were unable to find evidence for the proposed reversible carbonic anhydrase activity due to the uncoupling of the reaction by the mutation (**Table I**). A third alternative is a mechanism for the CupA/B active site involving a form of product trapping in which HCO_3_^-^, the hydration product of CO_2_, is tightly bound by its interaction with the nearby NdhF3/4-R37 immediately following its formation, thereby trapping the product. The trapped HCO_3_^-^ product is then proposed to be released by proton extraction due to proton-pumping by the membrane-spanning subunits (**Fig. S5**). According to this ‘product-trapping, energetic release’ hypothesis, the reaction starts with the normal nucleophilic attack by a Zn^2+^-hydroxyl anion upon incoming CO_2_, leading to a transient Zn-bicarbonate species. However, the HCO_3_^-^ displacement by H_2_O from its Zn^2+^ coordination position is then proposed to yield its rearrangement to bind between NdhF4-R37 and NdhF4-E114 through ionic and/or H-bonding interactions instead of releasing the product (**Figure S6, left panel)**. Computational calculations predict that the pK_a_s of NdhF4-R37 and NdhF4-E114 are very basic (pK_a_s ∼13), making such a docking of the HCO_3_^-^ electrostatically feasible [34]. A key aspect of this mechanism is the subsequent role of proton pumping by the NdhF and NdhD4 transmembrane subunits. Proton-pumping is theorized to remove a proton from the transient arginine-bicarbonate ionic adduct, thereby neutralizing its positive charge and diminishing its binding interaction, allowing for the release of bicarbonate. This model suggests a regulated bicarbonate production and release mechanism, intricately linking enzymatic activity to the electrochemical gradient. However, the feasibility of such a mechanism, including the strength of the ionic interaction of the HCO_3_^-^ between the NdhF4-R37 and NdhF4-E114 pair and the precise coordination chemistry of zinc, poses several challenges and would require rigorous experimental validation involving additional mutagenesis, kinetic studies, and structural-computational analysis.

## Supporting information

Paper_plus_Supplemental_Material

## 5. Acknowledgments

The authors would like to thank Dr. Zhifen Zhang for his assistance and advice on experiments and for the critical comments on the manuscript. We also thank Professor Dean Price for the strains at the foundation of this work and for the scientific advice over the course of this work. This work was supported by the United States Department of Energy, Office of Basic Energy Sciences, DE-FG02-08ER15968.

## Abbreviations

CA: carbonic anhydrase
CCM: CO_2_-concentrating mechanism
Cup: CO_2_ uptake proteins;
C_i_: inorganic carbon, primarily [HCO_3_^-^+CO_2_]
NDH-1_1-4_: Forms of the Type-1 NAD(P)H dehydrogenase complexes with NDH-1_3_ and NDH-1_4_ catalyzing CO_2_ hydration
CBB: Calvin-Bassham-Benson cycle of photosynthetic carbon fixation
CNDH-1: Type-1 proton-pumping dehydrogenase
RuBP: ribulose bisphosphate
Rubisco: ribulose bisphosphate carboxylase/oxygenase

## Footnotes

1 A mutant variant of Human Carbonic Anhydrase I has been crystallographically determined and shown to contain an arginine substitution of histidine 67 in the active site, but in this case, neither wild-type histidine 67 nor the substituting arginine coordinate the Zn ion responsible for anhydrase activity, but instead the arginine variant forms a new binding site for a second Zn ion that has esterase activity [37] M. Ferraroni, S. Tilli, F. Briganti, W.R. Chegwidden, C.T. Supuran, K.E. Wiebauer, R.E. Tashian, A. Scozzafava, Crystal Structure of a Zinc-Activated Variant of Human Carbonic Anhydrase I, CA I Michigan 1: Evidence for a Second Zinc Binding Site Involving Arginine Coordination, Biochemistry, 41 (2002) 6237-6244..

## Supplementary Materials

**Figure S1.**
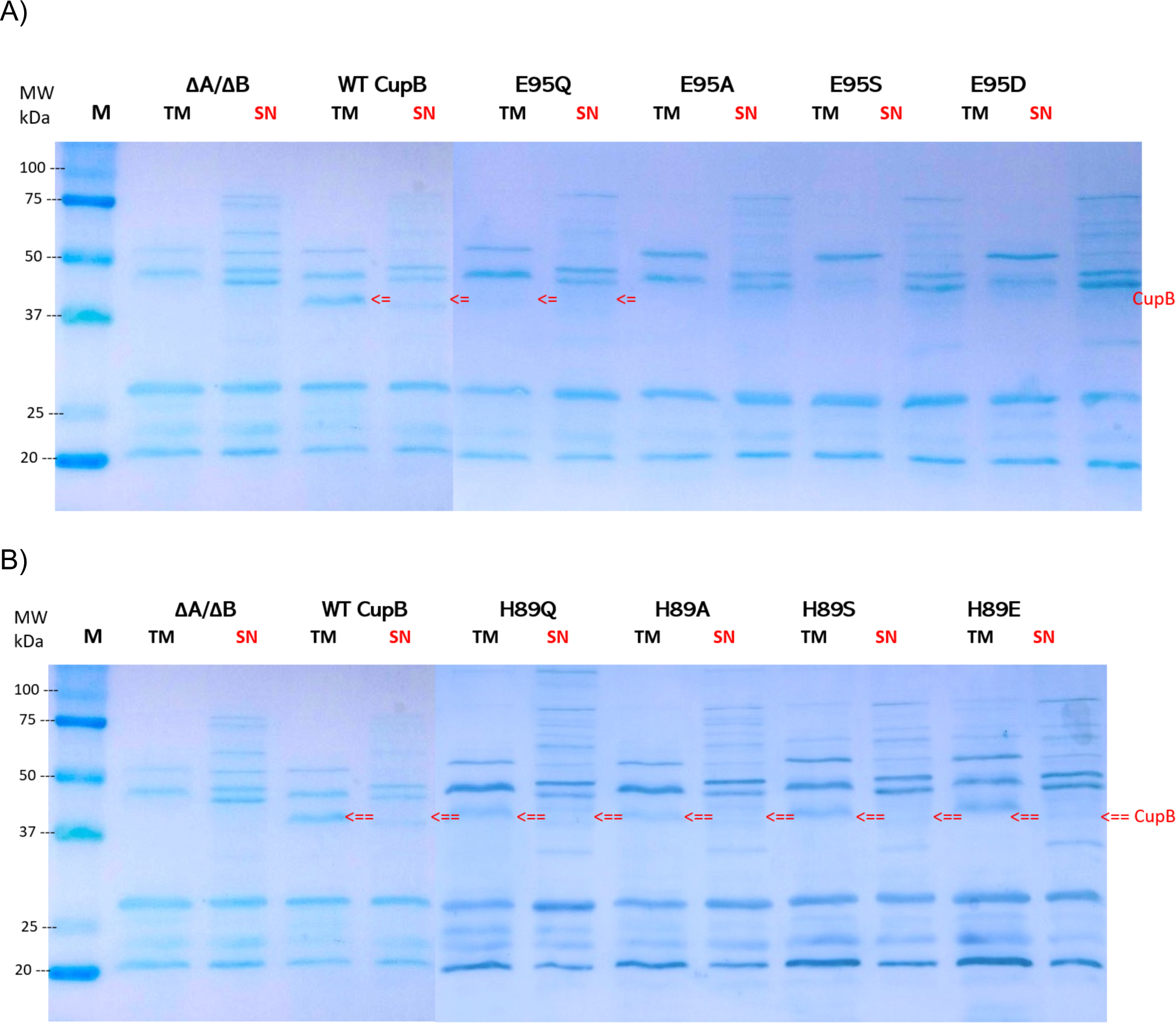
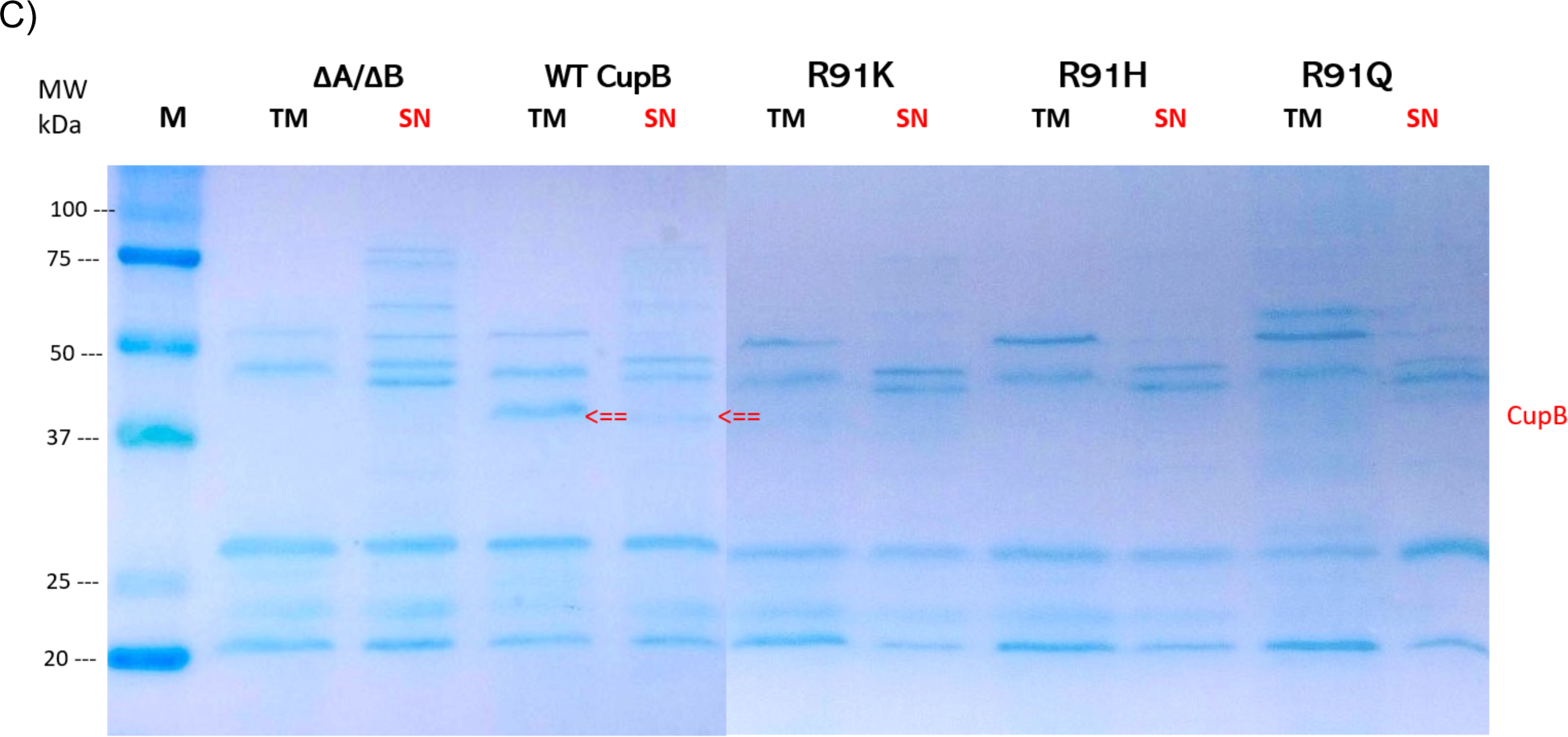
Membrane protein isolates of Synechococcus elongatus PCC sp. 7942 R91-CupB point mutants. Cells were grown at 3% CO_2_ in BG-11 medium at pH 8.0 and thylakoid membranes (TM) and soluble cytoplasmic fractions (SN) were centrifugally separated as described in Methods. After solubilization, samples were analyzed by SDS-PAGE, electroblotted to a polyvinylidene difluoride (PVDF) membrane, and probed with anti-CupB antibody. The expected molecular mass of CupB is 42 kDa. Panel A: CupB-Glu95 point mutants; Panel B: CupB-His89 point mutants; Panel C: CupB-Arg91 point mutants.

**Figure S2.**
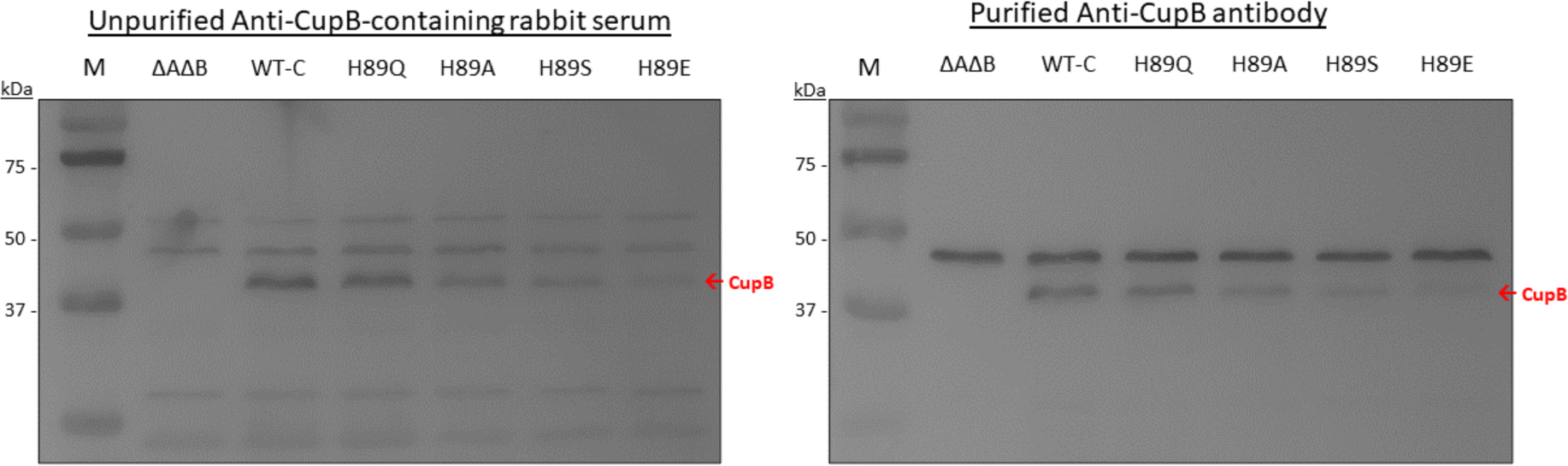
Purification of CupB antibody from rabbit serum. Thylakoid membrane proteins of *Syn*7942 CupB mutants are probed with CupB antibody present in the original rabbit serum (left panel) or CupB antibody purified from the rabbit serum using an immobilized synthetic peptide (right panel). Affinity purification was performed using the originally synthesized antigen (CupB peptide) as a ligand attached to a column matrix. Cyanobacterial cells were grown at 3% CO_2_ in BG-11 medium at pH 8.0 and thylakoid membranes were isolated as described in Methods. After solubilization, samples were analyzed by SDS-PAGE, electroblotted to a polyvinylidene difluoride (PVDF) membrane, and probed with anti-CupB immune serum (left panel) and affinity purified anti-CupB antibodies (right panel). A strong non-specific reaction is observed with an unknown polypeptide migrating at ∼47kDa in both the raw serum and affinity purified samples. CupB has an expected mass of 42 kDa.

### Possible mechanisms of CO_2_ hydration in NDH-1_3/4_ complexes

### Proton Product Removal Mechanism

The ‘*Proton Product Removal*’ hypothesis (**Fig S4**), depicting the proton released during CO_2_ hydration being integrated into the proton pumping activity of the antiporter-like subunits of the NDH-1 complex [1]. The hypothesis posits that the hydration reaction is driven by the removal of the product proton (H^+^), a process that is energetically coupled with the charge-relay system transmitted through the antiporter-like subunits. This coupling is essential for aligning the CO_2_ hydration reaction with the bioenergetics of the NDH-1_3/4_ complexes, facilitated by the removal of the proton generated during substrate water hydrolysis at the metal catalytic center. In this process, zinc (Zn) plays a key role by ① enabling the deprotonation of substrate H_2_O to form a hydroxide ion capable of ② executing a nucleophilic attack on the incoming CO_2_, akin to carbonic anhydrases (CAs), leading to ③ the release of HCO_3_^-^. The subsequent proton (H+) is effectively diverted from the hydroxide through efficient pumping by the antiporter domains, which transfer the proton via an internal proton transfer (PT) pathway to a proton loading site (PLS). This site is gated, specifically designed to trap the proton and prevent its back-reaction with the deprotonated metal hydroxide, thereby driving forward the overall reaction process.

**Figure S3.**
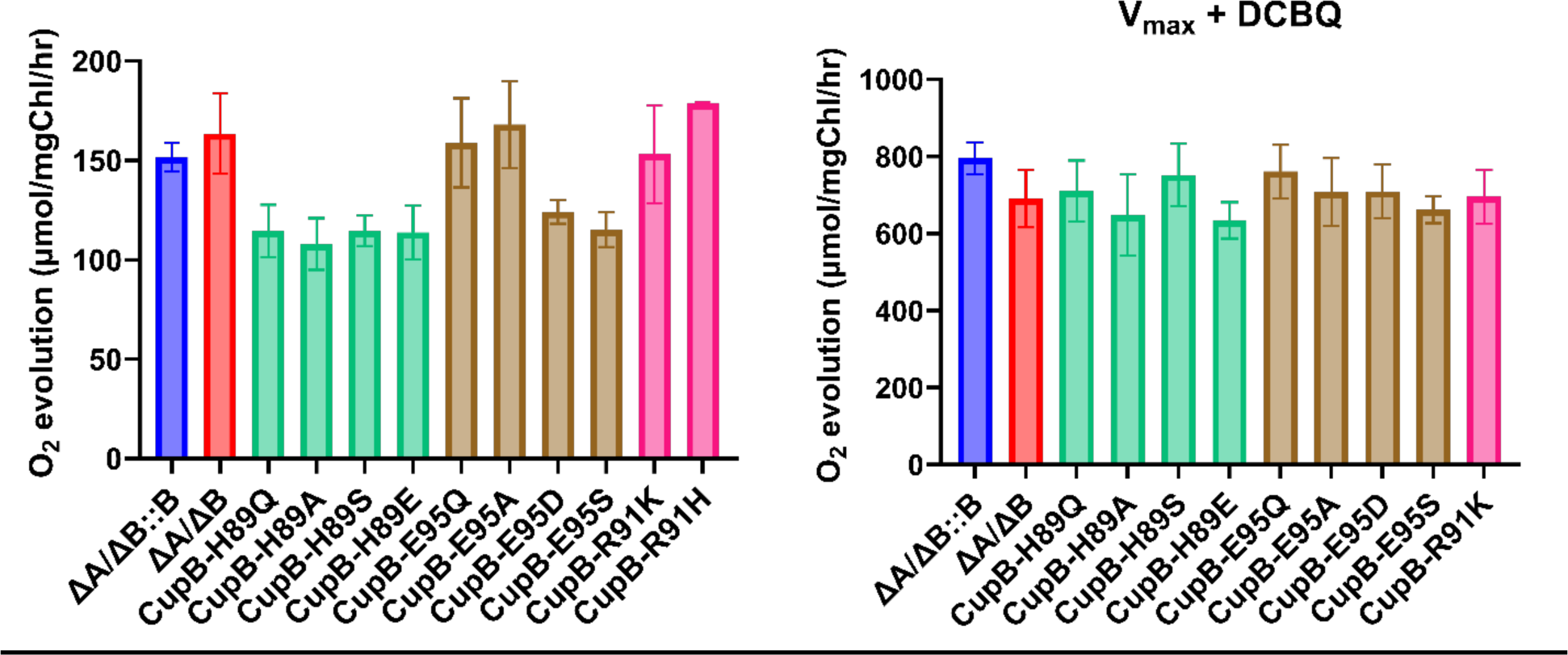
Clark-type electrode measurements of maximum O_2_ evolution coupled to CO_2_ fixation (left panel) and uncoupled using the artificial PSII electron acceptor, DCBQ (right panel). The CO_2_-dependent maximal rate corresponds to the maximum rate of O_2_ evolution measured during the 15-minute affinity assay (Fig. 4, main text) whereas the maximal, uncoupled rate of O_2_ evolution in the presence of DCBQ was obtained in the presence of 300 μM DCBQ and 1mM potassium ferricyanide.

### Product Trapping and Energetic Release Mechanism in CupA/B Active Site

This figure illustrates a proposed mechanism for the CupA/B active site, highlighting a form of product trapping where bicarbonate (HCO_3_^-^), the hydration product of CO_2_, is tightly bound to the NdhF3/4-R37 residue immediately after its formation. This interaction effectively traps the bicarbonate within the active site. The ‘product-trapping, energetic release’ hypothesis begins with a Zn^2+^-hydroxyl anion initiating a nucleophilic attack on incoming CO_2_, forming a transient Zn-bicarbonate species. Contrary to typical release, HCO_3_^-^ is then proposed to rearrange and bind between NdhF4-R37 and NdhF4-E114 through ionic or hydrogen bonding interactions instead of being released (**Fig. S5, left panel**). Computational predictions suggest that the pKas of NdhF4-R37 and NdhF4-E114 are highly basic (around pKa ∼13), supporting the feasibility of this bicarbonate docking through electrostatic interactions [2].

**Figure S4.**
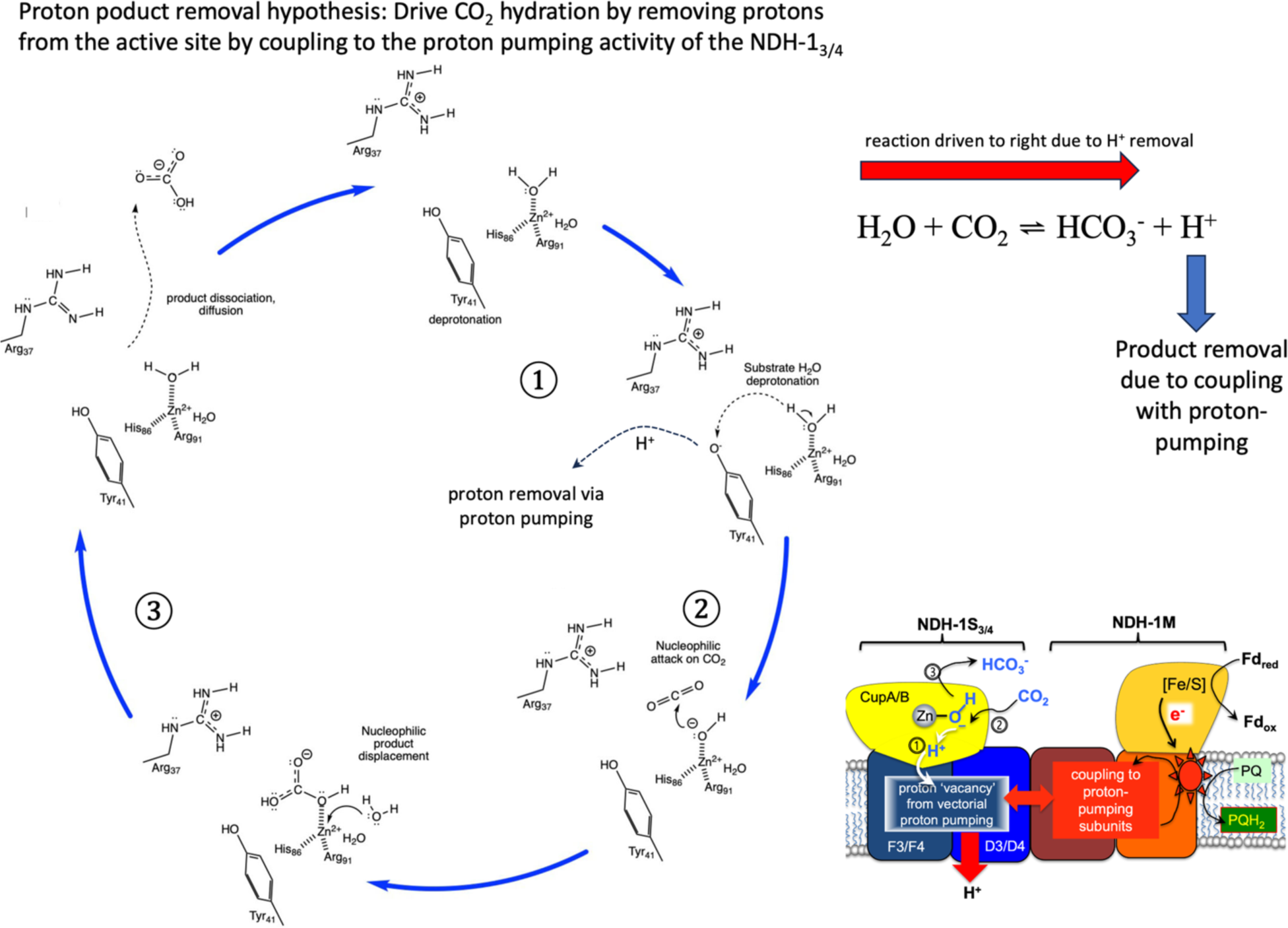

A critical aspect of this mechanism is the role of proton pumping by the NdhF and NdhD4 transmembrane subunits. It’s theorized that this proton-pumping activity extracts a proton from the arginine-bicarbonate adduct, neutralizing the positive charge of the arginine and weakening its interaction with bicarbonate, thereby facilitating bicarbonate release. This model implies a regulated bicarbonate production and release mechanism closely linked to the electrochemical gradient across the membrane. Nonetheless, the practicality of this mechanism, particularly the strength of the HCO_3_^-^ interaction with the NdhF4-R37 and NdhF4-E114 pair and the zinc coordination chemistry, presents challenges that necessitate thorough experimental validation, including mutagenesis, kinetic experiments, and structural-computational studies.

**Figure S5.**
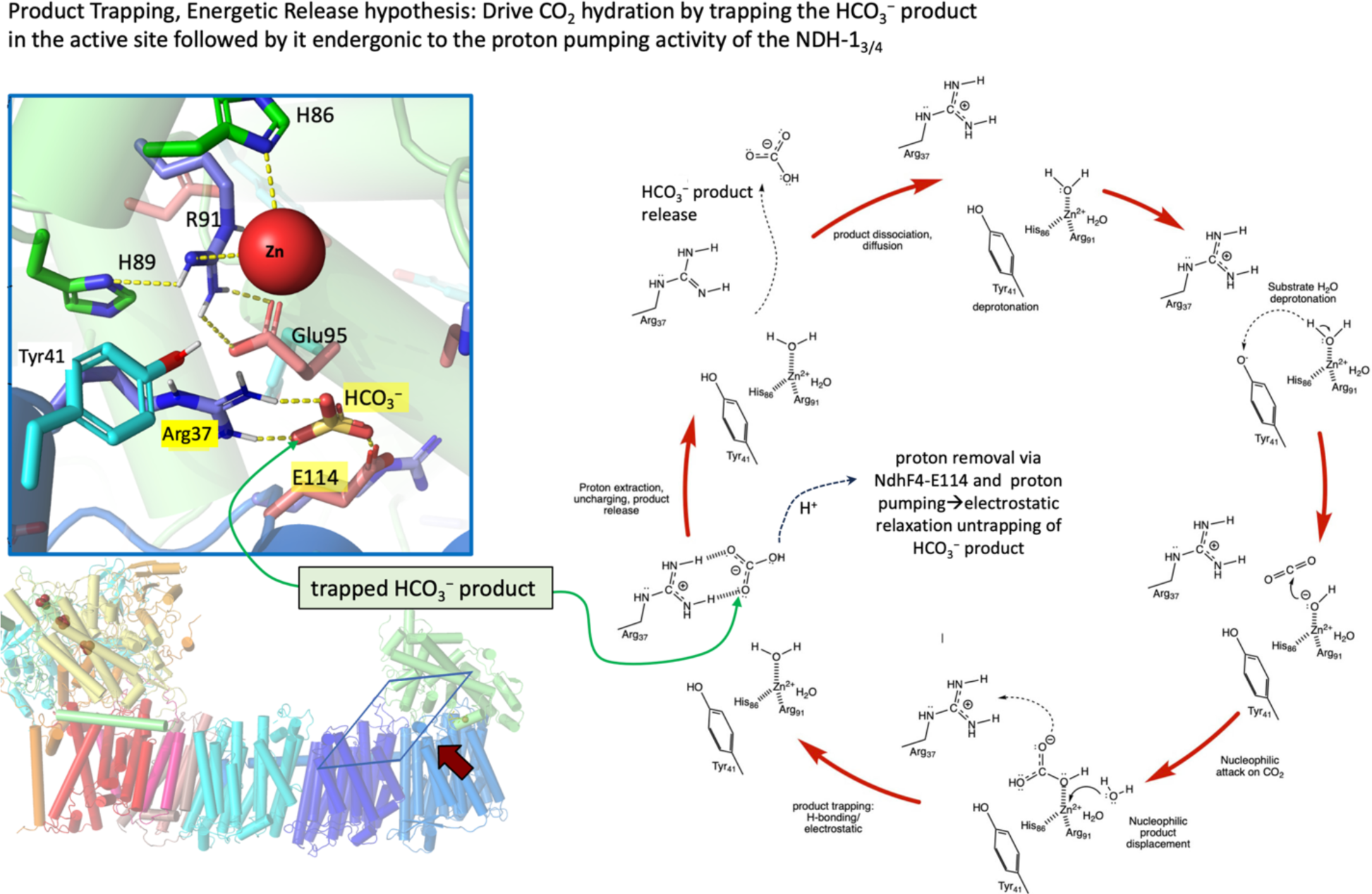

