## Supplementary material for "Elucidating the Role of Primary and Secondary Sphere Zn^2+^ Ligands in the Cyanobacterial CO_2_ Uptake Complex NDH-1_4_: The Essentiality of Arginine in Zinc Coordination and Catalysis": Paper_plus_Supplemental_Material

**Running head:** $CO_2$-concentrating mechanism

**Corresponding Author**
Robert L. Burnap
Oklahoma State University
307 Life Science East
Stillwater, OK 74078


**Research Category**
Photosynthesis, Membranes, and Bioenergetics

Title:

**Elucidating the Role of Primary and Secondary Sphere $Zn^{2+}$ Ligands in the Cyanobacterial $CO_2$ Uptake Complex NDH-$1_4$: The Essentiality of Arginine in Zinc Coordination and Catalysis**

Ross M. Walker, Minquan Zhang, and Robert L. Burnap
Department of Microbiology and Molecular Genetics, Oklahoma State University, Stillwater, OK 74078, USA

**Footnotes**

This work was supported by the U.S. Department of Energy Basic Energy Sciences; grant no. DE-FG02-08ER15968

**Key words**

carbonic anhydrase, CO2 concentrating mechanism; CCM; NDH-1; inorganic carbon, photosynthesis, zinc

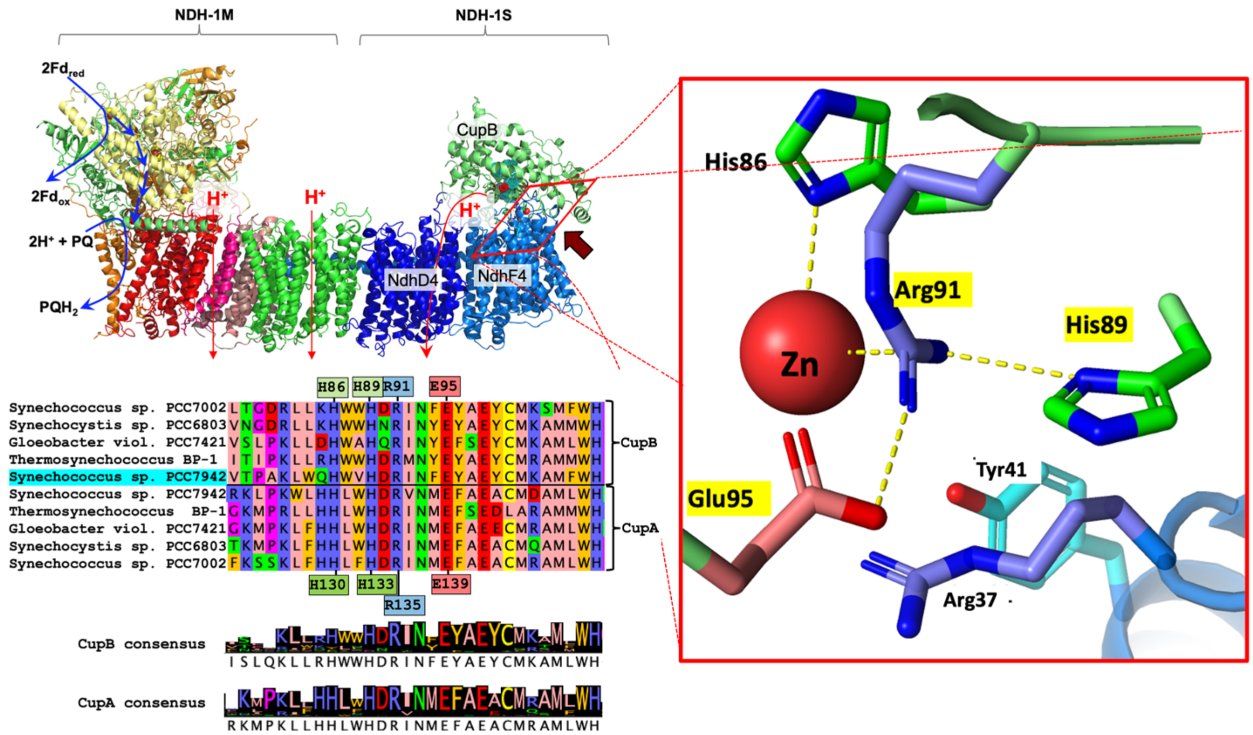


Figure 1. **The active site of $CO_2$ hydration. Multiple sequence alignment (MSA) of Cup proteins A and B from a relevant selection of cyanobacterial genomes (inset , lower left).** Correspondence between rows indicates homology according to MSA (ClustalW) analysis described previously [29]. Six species are shown as an excerpt from a larger alignment of CupA and CupB protein sequences identified from 157 cyanobacteria strains. The excerpted sequences correspond to *Synechococcus* sp. PCC7002 (Syn7002), *Synechocystis* sp. PCC6803 (*Syn*6803), *Gloeobacter violaceus* sp. PCC7421, *Thermosynechococcus elongatus* BP-1, *Synechococcus* sp. PCC7942 (*Syn*7942). The active site (the featured region of the sequences) shows high conservation as indicated by the consensus logos of the CupA and CupB protein subfamilies. **Homology modeling of the putative CupB active** site based on the CupA subunit of *Thermosynechococcus* from the NDH-$1_3$ published coordinates (PDB 6TJV) developed previously [29] as shown (**top left and right side, respectively)**. The mutationally targeted residues H89, R91, and E95 of CupB are indicated with yellow highlighted labels in the close-up of the Zn-containing active site (right side). The highly conserved residues around the putative active site of CupB and NdhF4 are shared by their respective homologs, CupA and NdhF3. The model shows two residues, CupB-H86 and CupB-R91, acting as direct ligands to the Zn ion, consistent with the homologous side chains of the CupA protein of the *Thermosynechococcus* NDH-$1_3$ structure.

---

[1] A mutant variant of Human Carbonic Anhydrase I has been crystallographically determined and shown to contain an arginine substitution of histidine 67 in the active site. In this case, neither wild-type histidine 67 nor the substituting arginine coordinate the Zn ion responsible for the primary anhydrase activity. However, the arginine variant forms a new binding site for a second Zn ion that has esterase activity, and possibly anhydrase activity [37] M. Ferraroni, S. Tilli, F. Briganti, W.R. Chegwidden, C.T. Supuran, K.E. Wiebauer, R.E. Tashian, A. Scozzafava, Crystal Structure of a Zinc-Activated Variant of Human Carbonic Anhydrase I, CA I Michigan 1: Evidence for a Second Zinc Binding Site Involving Arginine Coordination, Biochemistry, 41 (2002) 6237-6244..

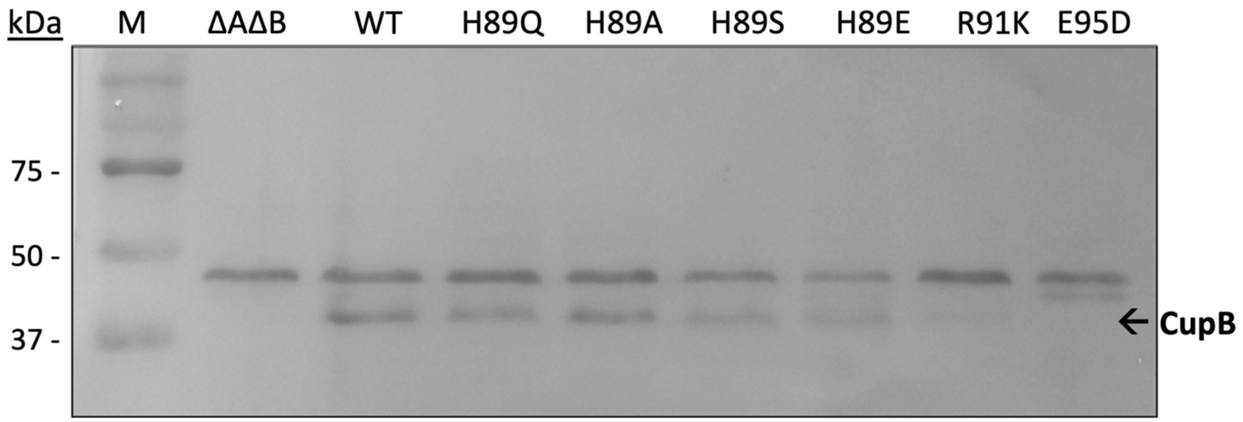


**Figure 2. Immunodetection of CupB accumulation in mutants *Syn*7942 strains.** Membrane protein isolates of *Syn*7942 strains lacking both CupA and CupB (ΔAΔB), expressing the wild-type CupB protein in the ΔAΔB genetic (WT-C) background and various CupB amino acid substitution mutants are probed with CupB antibody purified from rabbit immune serum. Cells were grown at 3% $CO_2$ in BG-11 medium at pH 8.0 and crude thylakoid membranes were isolated as described in Methods. Cytoplasmic fractions were also probed for the presence of CupB but none was detected (Fig S1). After solubilization, polypeptides in the membrane samples were analyzed by SDS-PAGE, electroblotted to a polyvinylidene difluoride (PVDF) membrane, and probed with anti-CupB antibody. CupB is expected to have a mass of 42 kDa.

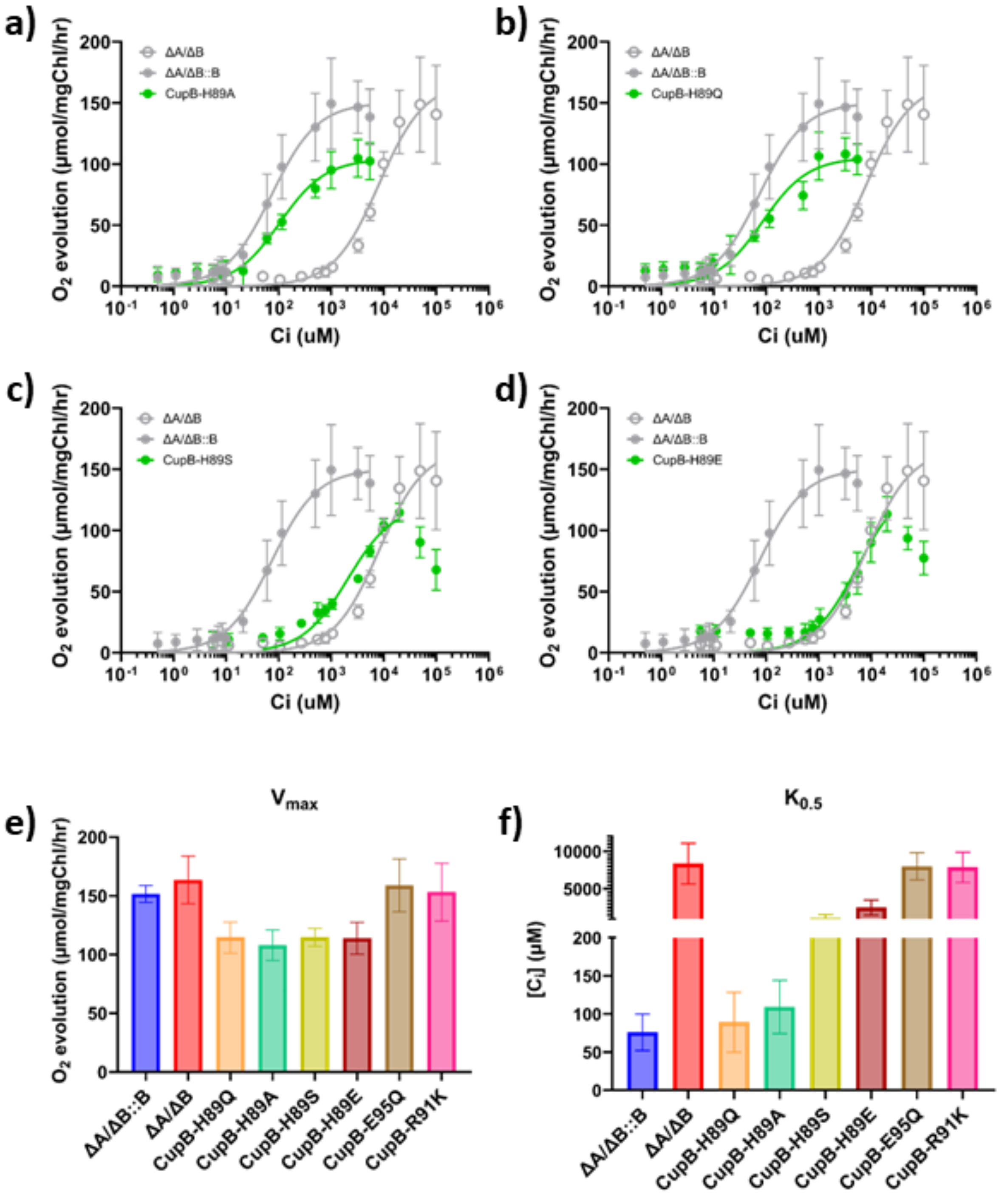


**Figure 3. Clark-type electrode measurements of $O_2$ evolution as a proxy for $CO_2$ uptake.** Whole cell response to titration of $C_i$ reveals the affinity for $CO_2$ in His89 mutants as a function of [$C_i$] (a – d). Cells were grown under 5% supplemental $CO_2$ to prevent accumulation of low-$C_i$ inducible CCM components such as SbtA or NDH-$1_3$. Affinity is measured as the [$C_i$] at which the cells evolve $O_2$ at half their maximum velocity. In this way, the affinity of whole cell

mutants can be represented by $K_{0.5}$, similar to the Michaelis-Menten $K_m$ unit. n = >3. $V_{max}$ BTP (e) represents the maximum rate of $O_2$ evolution measured during the 15-minute affinity assay. $K_{0.5}$ (f) represents the value of [$C_i$] which corresponds to half the $V_{max}$. Error bars represent standard deviation from the mean (biological n ≥ 3; technical n ≥ 8).

#### *3.4. No evidence for a mutationally-induced reversible carbonic anhydrase*

We had previously hypothesized that CupB-H89Q mutation uncoupled the proton pumping activity, rendering the Zn-centered carbonic anhydrase activity reversible and thus dissipating rather than promoting the formation of high bicarbonate concentrations in the cytoplasm [29]. To investigate this, we conducted carbonic anhydrase activity assays, measuring pH changes in diluted buffers with initially containing saturating concentrations of dissolved $CO_2$. Compared to the control using bovine carbonic anhydrase, anhydrase activity in membranes of the WT-control and the CupB-H89Q mutant was very low and, importantly, were equivalent. Thus, this test showed no significant differences in activity between the WT-control and the CupB-H89Q mutant (**Table 1**).

| Sample | Time (avg seconds): pH 8.3 → 6.3 | W-A units: $(T_0 - T)/T$ |
|---|---|---|
| Blank (water) | 145 ± 14 | nd |
| Bovine carbonic Anhydrase CII: | 22 ± 3 | 5.6 (expected 6.0) |
| WT-C isolated TMs | 151 ± 18 | nd |
| H89Q isolated TMs | 150 ± 16 | nd |

**Table 1. Carbonic anhydrase (CA) activity measured as the rate of change in pH of a weakly buffered solution from 8.3 to 6.3.** One WAU is defined as $(T_0 - T)/T$, where $T_0$ (uncatalyzed reaction) and $T$ (catalyzed reaction) are recorded as the time (s). Protocol adapted from Millipore Sigma (EC 4.2.1.1).

Ross M. Walker, Minquan Zhang, and Robert L. Burnap

Department of Microbiology and Molecular Genetics, Oklahoma State University, Stillwater, OK 74078, USA

**Figure S1.**

A)

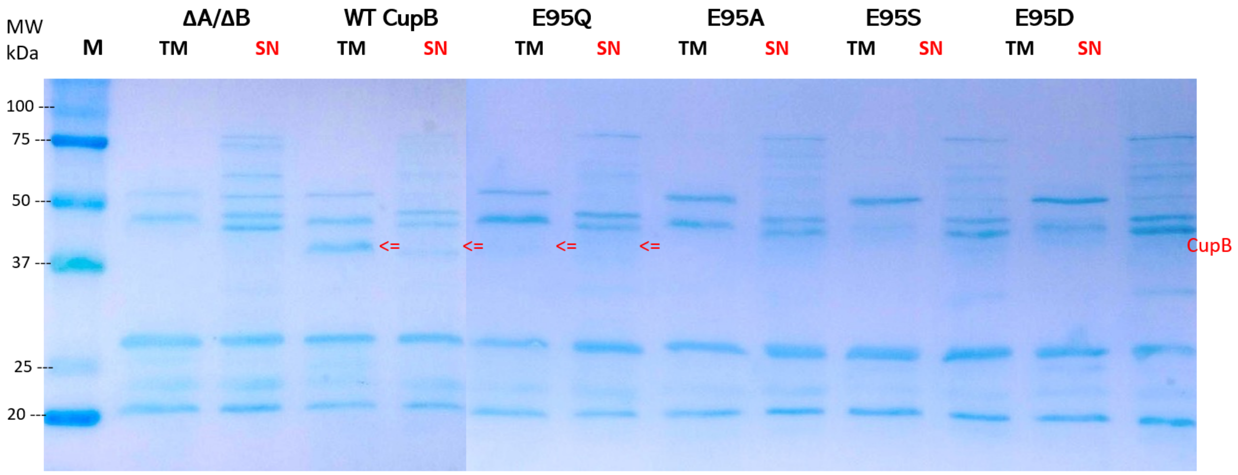
MW kDa
M
ΔA/ΔB
WT CupB
E95Q
E95A
E95S
E95D
TM SN TM SN TM SN TM SN TM SN TM SN
100
75
50
37
25
20
CupB

B)

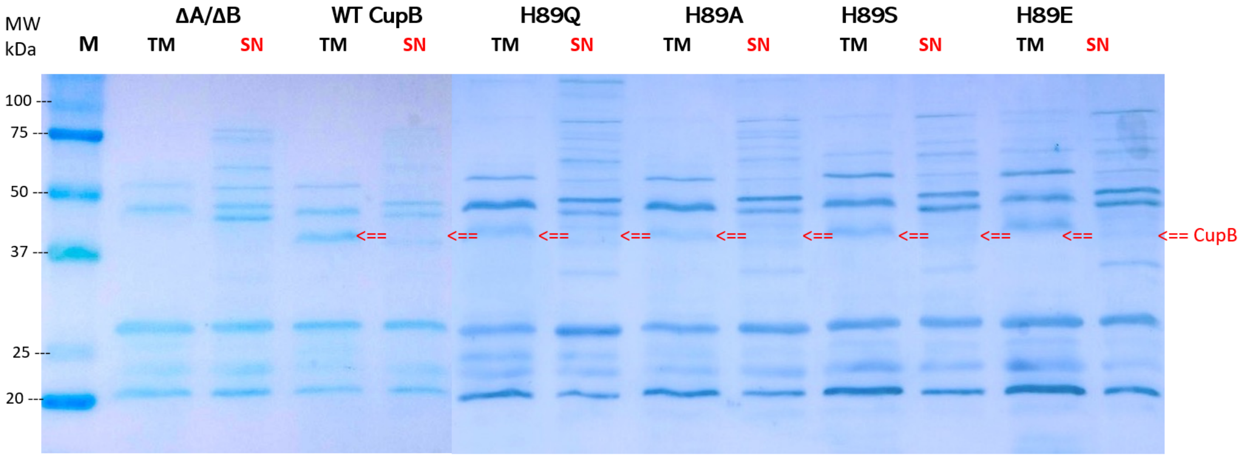
MW kDa
M
ΔA/ΔB
WT CupB
H89Q
H89A
H89S
H89E
TM SN TM SN TM SN TM SN TM SN TM SN
100
75
50
37
25
20
CupB

C)

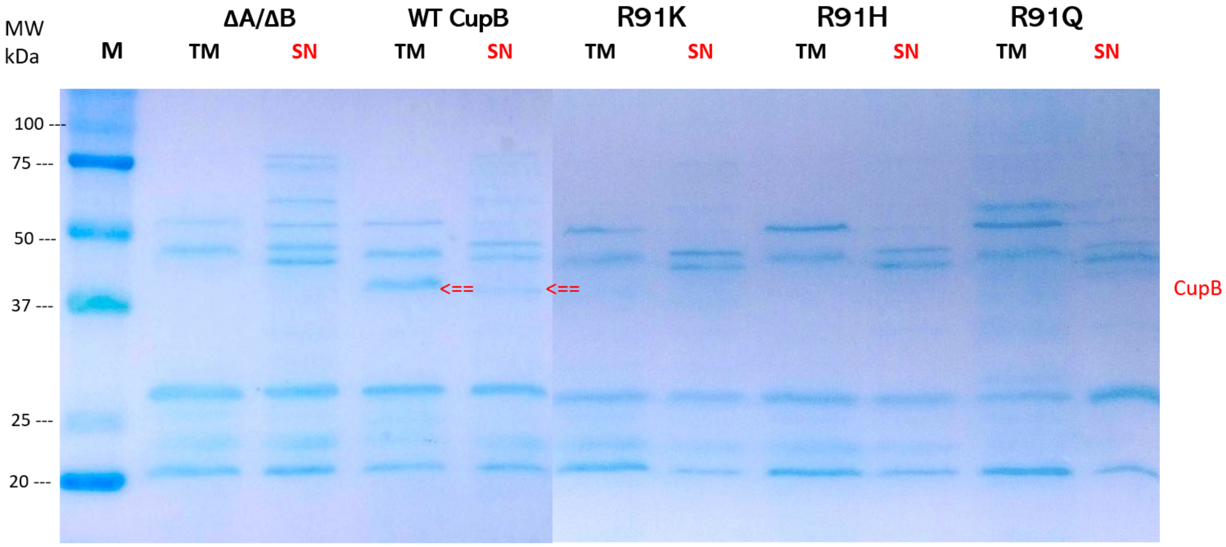


**Figure S1.** Membrane protein isolates of Synechococcus elongatus PCC sp. 7942 R91-CupB point mutants. Cells were grown at 3% $CO_2$ in BG-11 medium at pH 8.0 and thylakoid membranes (TM) and soluble cytoplasmic fractions (SN) were centrifugally separated as described in Methods. After solubilization, samples were analyzed by SDS-PAGE, electroblotted to a polyvinylidene difluoride (PVDF) membrane, and probed with anti-CupB antibody. The expected molecular mass of CupB is 42 kDa. Panel A: CupB-Glu95 point mutants; Panel B: CupB-His89 point mutants; Panel C: CupB-Arg91 point mutants.

### Figure S2

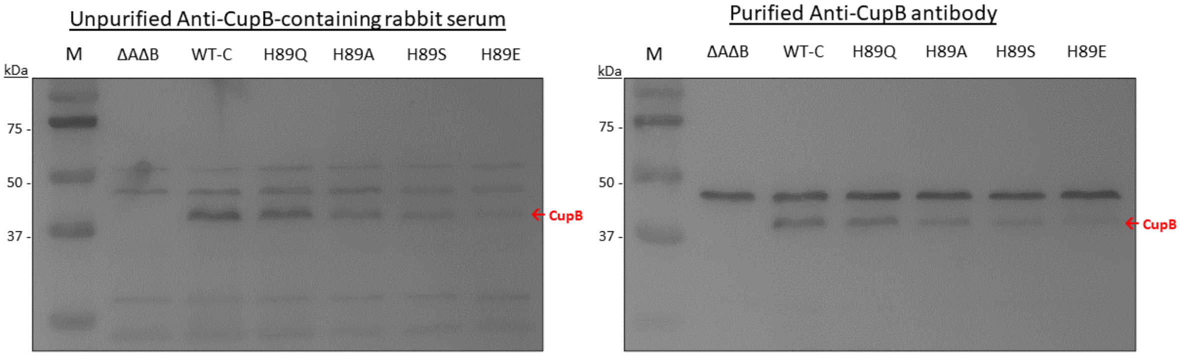


**Figure S2. Purification of CupB antibody from rabbit serum.** Thylakoid membrane proteins of *Syn*7942 CupB mutants are probed with CupB antibody present in the original rabbit serum (left panel) or CupB antibody purified from the rabbit serum using an immobilized synthetic peptide (right panel). Affinity purification was performed using the originally synthesized antigen (CupB peptide) as a ligand attached to a column matrix. Cyanobacterial cells were grown at 3% $CO_2$ in BG-11 medium at pH 8.0 and thylakoid membranes were isolated as described in Methods. After solubilization, samples were analyzed by SDS-PAGE, electroblotted to a polyvinylidene difluoride (PVDF) membrane, and probed with anti-CupB immune serum (left panel) and affinity purified anti-CupB antibodies (right panel). A strong non-specific reaction is observed with an unknown polypeptide migrating at ~47kDa in both the raw serum and affinity purified samples. CupB has an expected mass of 42 kDa.

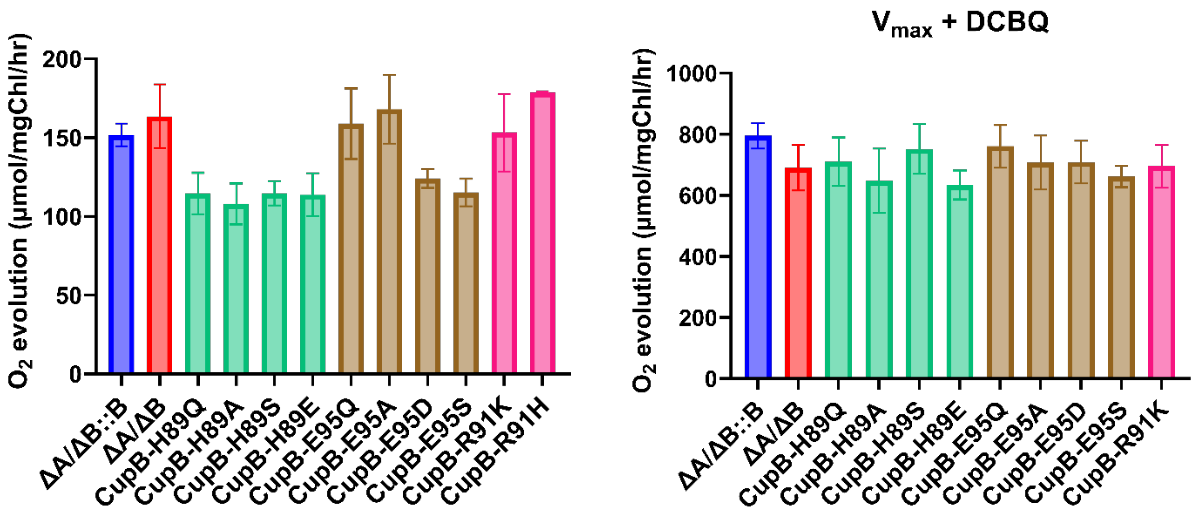


**Figure S3. Clark-type electrode measurements of maximum $O_2$ evolution coupled to $CO_2$ fixation (left panel) and uncoupled using the artificial PSII electron acceptor, DCBQ (right panel).** The $CO_2$-dependent maximal rate corresponds to the maximum rate of $O_2$ evolution measured during the 15-minute affinity assay (Fig. 4, main text) whereas the maximal, uncoupled rate of $O_2$ evolution in the presence of DCBQ was obtained in the presence of 300 µM DCBQ and 1mM potassium ferricyanide.

**Possible mechanisms of $CO_2$ hydration in NDH-$1_{3/4}$ complexes:**

**Figure S4**

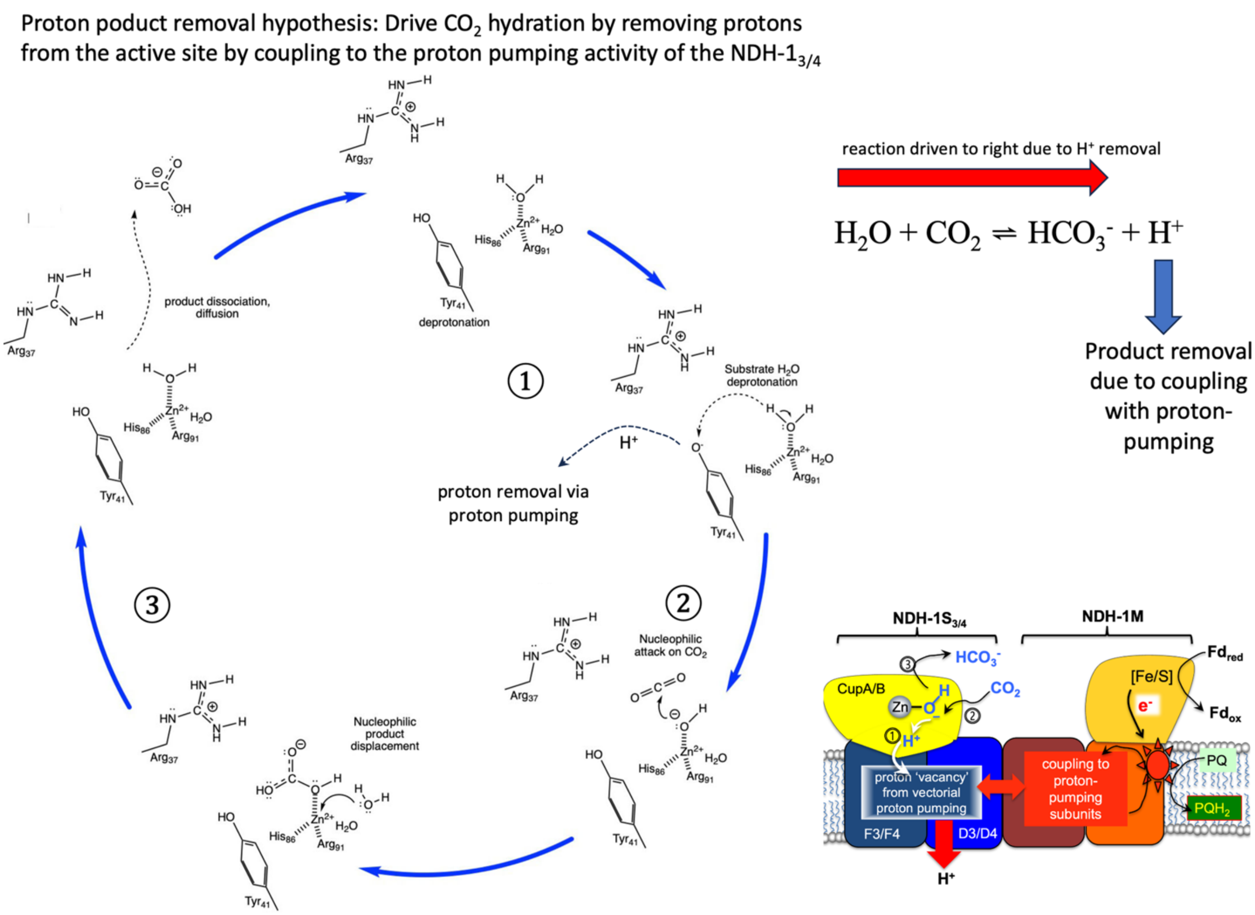


**Figure S4. The '*Proton Product Removal*' hypothesis**, depicting the proton released during $CO_2$ hydration being integrated into the proton pumping activity of the antiporter-like subunits of the NDH-1 complex [1]. The hypothesis posits that the hydration reaction is driven by the removal of the product proton ($H^+$), a process that is energetically coupled with the charge-relay system transmitted through the antiporter-like subunits. This coupling is essential for aligning the $CO_2$ hydration reaction with the bioenergetics of the NDH-$1_{3/4}$ complexes, facilitated by the removal of the proton generated during substrate water hydrolysis at the metal catalytic center. In this process, zinc (Zn) plays a key role by ① enabling the deprotonation of substrate $H_2O$ to form a hydroxide ion capable of ② executing a nucleophilic attack on the incoming $CO_2$, akin to carbonic anhydrases (CAs), leading to ③ the release of $HCO_3^-$. The subsequent proton (H+) is effectively diverted from the hydroxide through efficient pumping by the antiporter domains, which transfer the proton via an internal proton transfer (PT) pathway to a proton loading site (PLS). This site is gated, specifically designed to trap the proton and prevent its back-reaction with the deprotonated metal hydroxide, thereby driving the overall reaction process forward.

**Figure S5.**

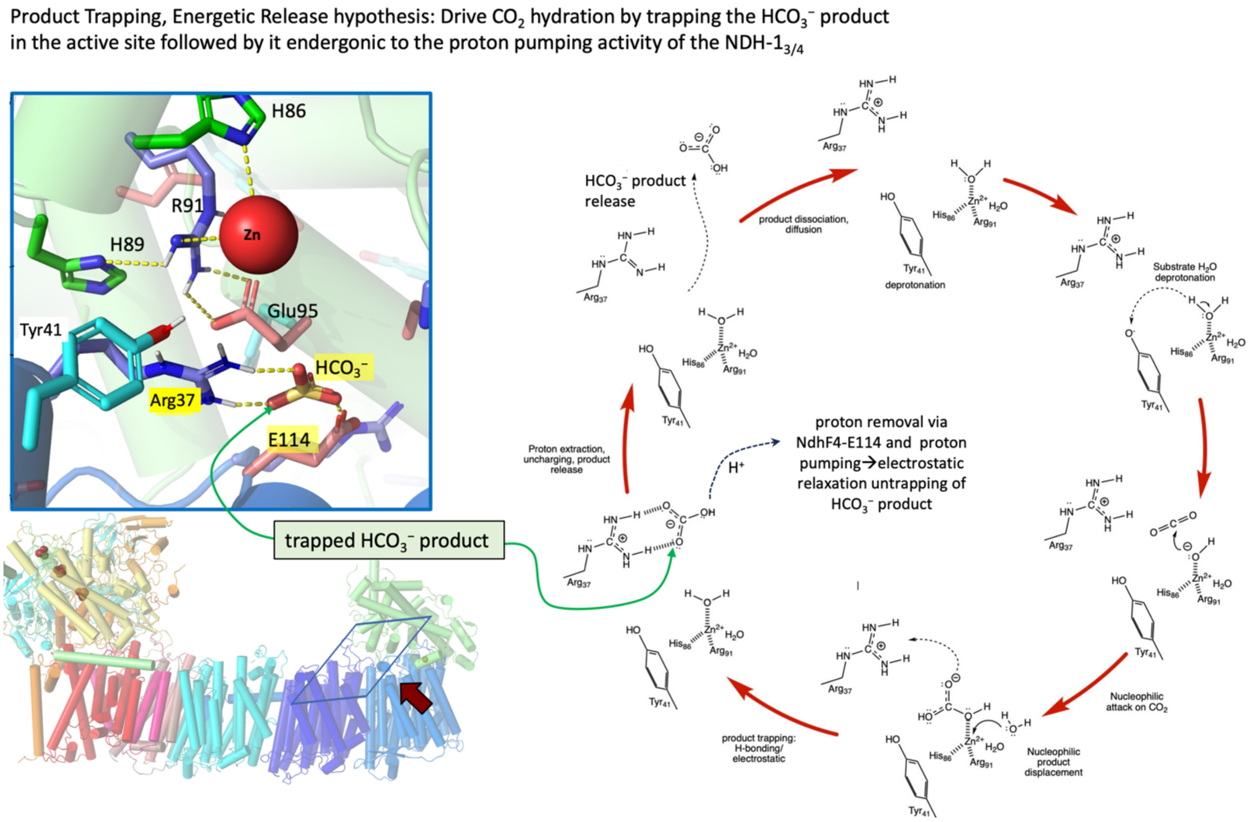


**Figure S5. Product Trapping and Energetic Release** This figure illustrates a proposed 'product trapping, energetic release' hypothesis. The CupA/B active site, highlighting a proposed product trapping where bicarbonate ($HCO_3^-$), the hydration product of $CO_2$, is tightly bound to the NdhF3/4-R37 residue and NdhF3/4-E114 immediately after its formation (left inset). This interaction effectively traps the bicarbonate within the active site. The 'product-trapping, energetic release' hypothesis begins with a $Zn^{2+}$-hydroxyl anion initiating a nucleophilic attack on incoming $CO_2$, forming a transient Zn-bicarbonate species. Contrary to typical release, $HCO_3^-$ is then proposed to rearrange and bind between NdhF4-R37 and NdhF4-E114 through ionic or hydrogen bonding interactions instead of being released (**Fig. S5, left panel**). Computational predictions suggest that the $pK_a$s of NdhF4-R37 and NdhF4-E114 are highly basic (around pKa ~13), supporting the feasibility of this bicarbonate docking through electrostatic interactions [2]

A critical aspect of this mechanism is the role of proton pumping by the NdhF and NdhD4 transmembrane subunits. It's theorized that this proton-pumping activity extracts a proton from the arginine-bicarbonate adduct, neutralizing the positive charge of the arginine and weakening its interaction with bicarbonate, thereby facilitating bicarbonate release. This model implies a regulated bicarbonate production and release mechanism closely linked to the electrochemical gradient across the membrane. Nonetheless, the practicality of this mechanism, particularly the strength of the $HCO_3^-$ interaction with the NdhF4-R37 and NdhF4-E114 pair and the zinc coordination chemistry, presents challenges that necessitate thorough experimental validation, including mutagenesis, kinetic experiments, and structural-computational studies.

A critical aspect of this mechanism is the role of proton pumping by the NdhF and NdhD4 transmembrane subunits. It's theorized that this proton-pumping activity extracts a proton from the arginine-bicarbonate adduct, neutralizing the positive charge of the arginine and weakening its interaction with bicarbonate, thereby facilitating bicarbonate release. This model implies a regulated bicarbonate production and release mechanism closely linked to the electrochemical gradient across the membrane.

### Supplemental Materials References

[1] J. Artier, R.M. Walker, N.T. Miller, M. Zhang, G.D. Price, R.L. Burnap, Modeling and mutagenesis of amino acid residues critical for CO2 hydration by specialized NDH-1 complexes in cyanobacteria, Biochimica et Biophysica Acta (BBA) - Bioenergetics, 1863 (2022) 148503.
[2] J.M. Schuller, P. Saura, J. Thiemann, S.K. Schuller, A.P. Gamiz-Hernandez, G. Kurisu, M.M. Nowaczyk, V.R.I. Kaila, Redox-coupled proton pumping drives carbon concentration in the photosynthetic complex I, Nat Commun, 11 (2020) 494.
